# CaSR modulates sodium channel-mediated Ca^2+^-dependent excitability

**DOI:** 10.1101/2021.03.03.433701

**Authors:** Briana J. Martiszus, Timur Tsintsadze, Wenhan Chang, Stephen M. Smith

## Abstract

Increasing extracellular [Ca^2+^] ([Ca^2+^]_o_) strongly decreases intrinsic excitability in neurons but the mechanism is unclear. By one hypothesis, [Ca^2+^]_o_ screens surface charge reducing voltage-dependent sodium channel (VGSC) activation and by another [Ca^2+^]_o_ activates Calcium-sensing receptor (CaSR) closing the sodium-leak channel (NALCN). Here we report that action potential (AP) firing rates increased in wild-type (WT), but not CaSR null mutant (Casr-/-) neocortical neurons, following the switch from physiological to reduced Ca^2+^-containing Tyrode. However, after membrane potential correction, AP firing increased similarly in both genotypes inconsistent with CaSR regulation of NALCN. Activation of VGSCs was the dominant contributor to the increase in excitability after the [Ca^2+^]_o_ change. VGSC conductance-voltage relationships were hyperpolarized by decreasing [Ca^2+^]_o_ for Casr-/- neurons indicating CaSR contributes to [Ca^2+^]_o_-dependent excitability via VGSCs. Regulation of VGSC gating by [Ca^2+^]_o_ is the key mechanism mediating [Ca^2+^]_o_-dependent changes in neocortical neuron excitability and CaSR influences neuronal excitability by its effects on VGSC gating.

## Introduction

Excitable tissues are strongly regulated by extracellular [Ca^2+^] ([Ca^2+^]_o_) (Neher and Sakaba, 2008; Ma et al., 2014; Jackman and Regehr, 2017). Movement of extracellular Ca^2+^, through voltage-activated Ca^2+^ channels (VACC), to the intracellular space is central to many of these processes (Nanou and Catterall, 2018). However, a distinct, extracellular mechanism that is independent of synaptic transmission also contributes to [Ca^2+^]_o_-dependent regulation of nerve and muscle function (Weidmann, 1955; Frankenhaeuser, 1957; Frankenhaeuser and Hodgkin, 1957). Decreases in [Ca^2+^]_o_ substantially facilitates action potential generation increasing intrinsic excitability(Weidmann, 1955; Frankenhaeuser, 1957; Frankenhaeuser and Hodgkin, 1957). In the brain, physiological neuronal activity decreases [Ca^2+^]_o_ (Nicholson et al., 1978) leading to further increases in action potential firing in neighboring neurons (Anderson et al., 2013). The firing patterns and computational properties of local circuits are impacted substantially by this positive feedback leading to changes in brain behaviors (Titley et al., 2019). Furthermore under pathological conditions, larger decreases in [Ca^2+^]_o_ occur, resulting in even greater changes in circuit activity, and implicating [Ca^2+^]_o_-dependent excitability in the pathogenesis of brain injury (Ayata and Lauritzen, 2015).

Classical studies proposed that the mechanism underlying [Ca^2+^]_o_-dependent excitability centers on voltage-gated sodium channel (VGSC) sensitivity to extracellular Ca^2+^. External Ca^2+^ was proposed to interact with local negative charges on the extracellular face of the membrane or ion channels thereby increasing the potential field experienced by VGSCs and reducing the likelihood of VGSC activation at the resting membrane potential (Frankenhaeuser and Hodgkin, 1957; Hille, 1968). Reductions in [Ca^2+^]_o_ decrease the screening of the surface potential, reverse the membrane stabilization and facilitate VGSC activation (Frankenhaeuser and Hodgkin, 1957; Hille, 1968). This surface potential screening model accounted for [Ca^2+^]_o_-dependent excitability in nerves and muscle without a need for additional molecular players and was widely accepted (Hille, 2001). However, the theory was challenged by new data demonstrating that activation of the sodium leak channel (NALCN), a non-selective cation channel, by the intracellular proteins, UNC79 and UNC80 (Lu et al., 2009; Lu et al., 2010) was necessary for [Ca^2+^]_o_-dependent excitability to occur in hippocampal neurons. Following the deletion of NALCN or UNC79, [Ca^2+^]_o_-dependent excitability was completely lost suggesting the increased excitability resulted from the activation of the non-rectifying NALCN which depolarized neurons and increased the likelihood of action potential generation <u>independent</u> of changes in VGSC function (Lu et al., 2010). The calcium-sensing receptor (CaSR), a G-protein coupled receptor (GPCR), was hypothesized to detect and transduce the [Ca^2+^]_o_ changes and signal to the downstream multistep pathway(Lu et al., 2010). CaSR is well-positioned as a candidate [Ca^2+^]_o_ detector because at nerve terminals it detects [Ca^2+^]_o_ and regulates a non-selective cation channel(Smith et al., 2004; Chen et al., 2010) and because it transduces changes in [Ca^2+^]_o_ into NALCN activity following heterologous co-expression of CaSR, NALCN, UNC79, and UNC80 (Lu et al., 2010). Interest in the UNC79-UNC80-NALCN pathway has also risen, due to its essential role in the maintenance of respiration(Lu et al., 2007), the regulation of circadian rhythms (Lear et al., 2013; Flourakis et al., 2015), and because mutations of UNC80 and NALCN cause neurodevelopmental disorders, characterized by development delay and hypotonia (Al-Sayed et al., 2013; Perez et al., 2016).

Here we address the question of whether the G-protein mediated NALCN pathway or VGSCs transduce the [Ca^2+^]_o_-dependent effects on excitability. We test if CaSR is a modulator of neuronal excitability via its action on a nonselective cation channel, determine the impact of CaSR expression on factors of intrinsic neuronal excitability, and examine the relative contributions of [Ca^2+^]_o_-regulated changes on VGSC and NALCN gating. In recordings from neocortical neurons, isolated by pharmacological block of excitatory and inhibitory inputs, we determine that neuronal firing is increased by decreasing external divalent concentrations and that this is almost entirely attributable to [Ca^2+^]_o_-dependent shifts in VGSC gating. Surprisingly, CaSR deletion substantially shifted VGSC gating, but had no effect on NALCN sensitivity to [Ca^2+^]_o_. Taken together our experiments indicate that acute [Ca^2+^]_o_-dependent increases in neuronal excitability result from changes in VGSC and NALCN gating and that CaSR contributes by an, as yet, uncharacterized action on VGSCs.

## Results

### CaSR and [Ca^2+^]_o_-dependent neuronal excitability

The elimination of [Ca^2+^]_o_-dependent excitability in neurons by deletion of UNC79 or NALCN challenged the long-standing hypothesis that local or diffuse surface charge screening of VGSCs mediated these effects(Lu et al., 2010). But how were changes in external divalent ion concentrations transduced to UNC79 and NALCN? We tested if CaSR provided the link, by comparing neuronal excitability in wild-type (WT) and nestin cre-recombinase expressing CaSR null-mutant (creCasr^-/-^) neurons that were genotyped by PCR (see Methods)(Chang et al., 2008). Quantification by RT-qPCR indicated >98% reduction in the Casr expression levels in neocortical cultures produced from creCasr^-/-^ mice compared to cre-positive WT (creWT; Figure 1-supplement). Current clamp recordings were performed to measure the intrinsic spontaneous action potential firing rate from pharmacologically-isolated, cultured, neocortical neurons (glutamatergic and GABAergic activity blocked by 10 µM CNQX, 50 µM APV, and 10 µM Gabazine respectively). After establishing the whole-cell configuration, we measured the action potential firing rates of conventional WT (conWT), creWT, and creCasr^-/-^ neurons in physiological Tyrode solution (; containing 1.1 mM [Ca^2+^] and 1.1 mM [Mg^2+^]) at the resting membrane potential (RMP) and then in reduced Ca^2+^ and Mg^2+^ Tyrode (T_0.2_; containing 0.2 mM [Ca^2+^] and [Mg^2+^]). The reduction in [Ca^2+^]_o_ and [Mg^2+^]_o_ caused a small depolarization and an increase in spontaneous action potential firing in both WT (con- and cre-) neurons within 15 seconds of the solution change that was substantially attenuated in the creCasr^-/-^ neuron (Figure 1A, middle row). Changing back to physiological external divalent concentrations reversed this effect within 10 s (Figure 1A, lower row). The pooled data from similar experiments indicated that on average the conWT and creWT neurons were equally sensitive to decreased [Ca^2+^]_o_ and [Mg^2+^]_o_ and had similar basal levels of activity (Figure 1B-D). Two-way repeated measures (RM) ANOVA confirmed a significant interaction indicating the response to changes in [Ca^2+^]_o_ and [Mg^2+^]_o_ were dependent on genotype (F (2,54) = 3.193, *P* = 0.049, Supplementary Table 1). Post-hoc tests (Sidak compensated for multiple comparisons here and in all later tests) confirmed that the reduction in [Ca^2+^]_o_ and [Mg^2+^]_o_ substantially increased action potential frequency in conWT and creWT but not creCasr^-/-^ neurons (Figure 1B, P =0.0009, <0.0001, and =0.6697, respectively). These data indicate that CaSR loss substantially reduces [Ca^2+^]_o_-dependent excitability in neurons.

**Figure 1:**
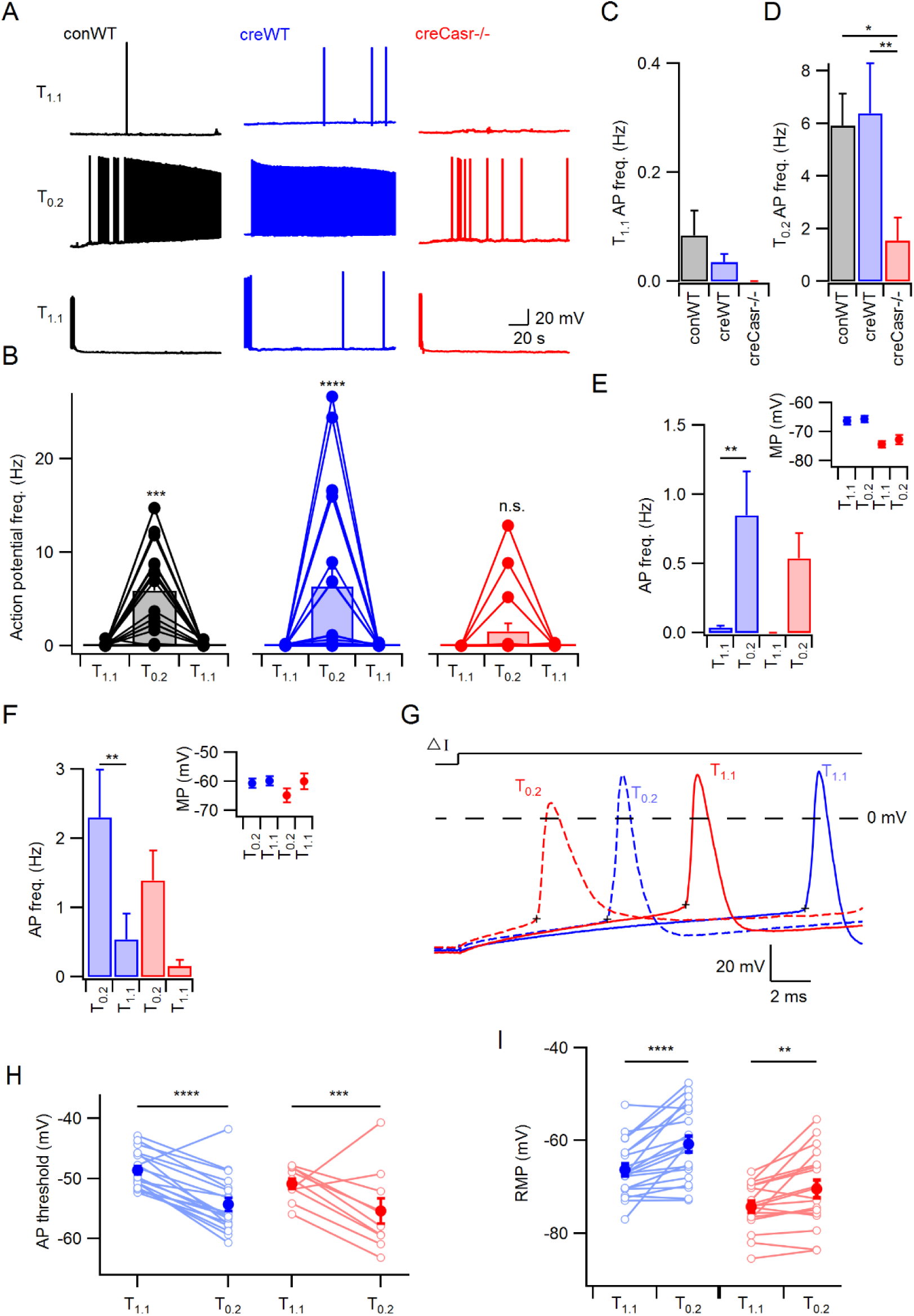
CaSR deletion reduces [Ca^2+^]_o_-dependent excitability. A) Spontaneous voltage traces at RMP following the application of solutions with different divalent concentrations (_1.1_ (upper traces), T_0.2_ (middle), and recovery (lower)) recorded in three individual neurons with or without CaSR (conWT (black), creWT (blue) and creCasr^-/-^ (red)). CreCasr^-/-^Each trace depicts 150 s of continuous acquisition. B) Histograms of average action potential (AP) frequency (Hz) recorded using the same solutions:, T_0.2_, and recovery. Individual recordings represented by open circles linked with lines and average is represented with a bar. From left to right graphs depict conWT (n = 18), creWT (n = 21), and CreCasr^-/-^ (n = 18). ANOVA: Post-hoc tests (Sidak compensated for multiple comparisons here and in all later figures) showed that AP frequency increased in conWT (P = 0.0009) and cre WT (P, 0.0001), but not CaSr^-/-^ (P = 0.6697) neurons when changing from to T_0.2_. C) Baseline average action potential frequency in. was unaffected by genotype (P > 0.999) D) Average action potential frequency with T_0.2_ application was the same in conWT and creWT (P = 0.9831) and higher than in creCasr^-/-^ neurons (P = 0.013 and 0.0033 respectively). E) Histogram summarizing effects of [Ca^2+^]_o_ on AP frequency in creWT and Casr^-/-^ neurons after a current injection to counter [Ca^2+^]_o_-dependent depolarization following T_0.2_ application. Two-way RM ANOVA indicates that reducing [Ca^2+^]_o_ increases the AP frequency (F (1, 35) = 11.54, P = 0.0017) and that this is significant in the creWT but not creCasr^-/-^neurons (P = 0.0075 and 0.1555 for 21 and 16 recordings respectively). Inset shows average membrane potential after the current injection. F) Histogram summarizing effects of [Ca^2+^]_o_ on AP frequency in creWT and Casr^-/-^ neurons after current injection in to depolarize membrane potential to value recorded in T_0.2_. Two-way RM ANOVA indicates that reducing [Ca^2+^]_o_ increases the AP frequency (F (1, 35) = 45.09, P = 0.0004) and that this is significant in the creWT but not creCasr^-/-^ neurons (P = 0.0044 and 0.056 for 21 and 16 recordings respectively). Inset shows average membrane potential after the current injection. G) Exemplar action potentials elicited by current injection in a creWT (blue) and a creCasr^-/-^ neuon (red) in (unbroken) and T_0.2_ (broken). Action potential threshold is indicated by + for the first AP elicited by current injection (50 to 200 pA) under the same conditions as panel E. H) Plot of average AP threshold in and T_0.2_ in creWT and creCasr^-/-^ neurons elicited as per panel G. Two-way RM ANOVA indicates that reducing [Ca^2+^]_o_ hyperpolarized the AP threshold (F (1, 27) = 56.48, P < 0.0001) but that genotype had no effect (F (1, 27) = 2.284, P = 0.1424). [Ca^2+^]_o_ was highly effective in both creWT and creCasr^-/-^neurons (P < 0.0001 and P = 0.0003 for 19 and 10 recordings respectively). Individual neuron values are represented by open circles linked by lines and averages by filled circles. I) Plot of effect of [Ca^2+^]_o_ and CaSR on RMP. Two-way RM ANOVA indicates that increasing [Ca^2+^]_o_ (F (1, 37) = 31.65, P < 0.0001) and CaSR deletion (F (1, 37) = 19.1, P < 0.0001) hyperpolarized the RMP without an interaction (F (1, 37) = 1.035, P = 0.3155). Post-hoc testing indicated RMP was depolarized with the switch to T_0.2_ in both creWT and creCasr^-/-^neurons (P < 0.0001 and P = 0.0066 for 21 and 18 recordings respectively).

### CaSR-independent mechanisms contribute to [Ca^2+^]_o_-dependent signaling

If CaSR-regulated NALCN-dependent depolarization is entirely responsible for the extracellular divalent-sensitive changes in neuronal excitability (Lu et al., 2010) then CaSR signaling to NALCN would depolarize the membrane potential when extracellular calcium was reduced. The same hypothesis predicts that reversal of this depolarization should prevent (or block) the increase in excitability seen in T_0.2._ To test this prediction we measured action potential frequency following current injections to drive the membrane potential in low [Ca^2+^]_o_ conditions to match the RMP seen during physiological [Ca^2+^]_o_ conditions (unique to each neuron). Action potential frequency in T_0.2_ was reduced by the hyperpolarization but neurons remained sensitive to changes in [Ca^2+^]_o_ (Figure 1E, Suppl. Table 2; F (1,35)=11.54, P=0.0017, 2-way RM ANOVA) indicating mechanisms besides NALCN were involved. Post-hoc tests confirmed this retained sensitivity to [Ca^2+^]_o_ and [Mg^2+^]_o_ was restricted to creWT and not creCasr^-/--^ neurons, (Figure 1E; P=0.0075 and =0.156, respectively). Similarly, in the reciprocal experiment in which the membrane potential in was depolarized to match that at low divalent concentration, the decrease in external divalent concentration increased action potential frequency (Figure 1F, Suppl. Table 3; F (1,35)=15.17, P=0.0004, 2-way RM ANOVA), and this was significant in creWT but not creCasr^-/-^ neurons (Figure 1F, P = 0.004). Ineffective matching of the membrane potential following solution changes did not account for the persistence of [Ca^2+^]_o_-dependent excitability (insets, Figure 1E,F). These data indicate a mechanism in addition to CaSR-NALCN-mediated depolarization must contribute to the extracellular divalent-sensitive changes in neuronal excitability.

### Does CaSR modulate RMP and [Ca_2+_]_o_-dependent depolarization?

We tested if there was an endogenous difference in the excitability of creWT and creCasr^-/-^ neurons to explain the retained sensitivity to divalents by measuring the action potential threshold. Action potentials were elicited in and T_0.2_ using minimal current injection (50-250 pA) and the threshold measured as the point at which dV/dt reached 20 mV/ms (Figure 1G, membrane potential-corrected as in Figure 1 E). The action potential threshold was hyperpolarized from −48.6 ± 0.7 mV to −54.3 ± 1.1 mV with the switch from to T_0.2_ in creWT neurons (Figure 1H) which would have increased excitability. However, the same effect was observed in creCasr^-/-^ neurons (−50.9 ± 0.86 mV to −55.4 ± 2.1 mV; F (1,27)=56.48, P < 0.0001, 2-way RM ANOVA, Suppl. Table 4). The lack of effect of CaSR on AP threshold (F (1, 27) = 2.284, P = 0.142) in these experiments, indicated the reduced divalent sensitivity of creCasr^-/-^ (Figure 1E,F) was not simply due to altered action potential threshold.

We next asked if differences in RMP in and the response of RMP to T_0.2_ could contribute to the different divalent-dependent excitability of creWT and creCasr^-/-^ neurons. Genotype and external divalent concentrations were both significant determinants of RMP (zero current injection; 2-way RM ANOVA, (Suppl. Table 5, F (1,37)=19.1, P < 0.0001 and F (1,37)=31.65, P < 0.0001, respectively). As CaSR deletion did not affect AP threshold (Figure 1H), action potential generation presumably occurred more frequently in the creWT neurons due to the relatively depolarized membrane potential (8 mV positive than creCasr^-/-^ neurons, Figure 1I,).

If CaSR-mediated NALCN-dependent depolarization is sufficient to account for [Ca^2+^]_o_-dependent excitability then creWT but not creCasr^-/-^ neurons, should depolarize in response to decreased [Ca^2+^]_o_. However, the RMP of creWT and creCasr^-/-^ neurons both depolarized similarly (Figure 1I; 5.6 ± 1.1 mV, P < 0.0001 and 3.9 ± 1.2 mV, P = 0.0066 respectively) when T_0.2_ was applied indicating the existence of a [Ca^2+^]_o_-sensitive pathway in creCasr^-/-^ neurons. Overall these data support the idea that CaSR played a substantial role in mediating divalent dependent changes in excitability, but that WT neurons also possessed CaSR-independent mechanisms to fully account for the [Ca^2+^]_o_-dependent excitability.

### CaSR-independent mechanisms contribute to [Ca^2+^]_o_-dependent signaling

Further mechanistic complexity was suggested by the effects of CaSR and [Ca^2+^]_o_ on RMP. This lead to a number of additional questions including: does the difference in RMP contribute to the difference in [Ca^2+^]_o_-dependent excitability between creWT and creCasr^-/-^ neurons, how do decreases in [Ca^2+^]_o_ depolarize creCasr^-/-^ neurons, and is this pathway present in creWT neurons? To address the first of these questions, we compared the response of creWT and creCasr^-/-^ neurons to changes in extracellular divalent concentrations after removing the confounding variation in RMP. After establishing a stable current-clamp recording in we injected a standing current (I_a_) until the resting membrane potential was −70 mV. We then recorded for 50 s before switching the bath solution to T_0.2_. As before, there was a small depolarization followed by an increase in action potential frequency in creWT neurons (Figure 2A,B). To test if this increase in excitability was fully attributable to [Ca^2+^]_o_-dependent depolarization we adjusted the standing current (I_b_) until the membrane potential was −70 mV and then measured the action potential frequency (Figure 2C). In the exemplar, AP firing was reduced by the hyperpolarization but remained higher in T_0.2_ at −70 mV than in at −70 mV (Figure 2A-C) confirming CaSR-mediated depolarization was not acting alone to increase the excitability (c.f. Figure 1I). The creCasr^-/-^ neurons responded similarly to T_0.2_ and hyperpolarization (Figure 2A-C) indicating the effect was not mediated by CaSR. We compared the average effects of at −70 mV with I_a_, T_0.2_ with I_a_, and T_0.2_ at −70 mV with I_b_ on creWT and creCasr^-/-^ genotypes (Figure 2D, Suppl. Table 6) using a 2-way RM ANOVA. Extracellular divalent concentration and membrane potential substantially affected action potential frequency (F (3, 87) = 17.97, P < 0.0001). CaSR deletion did not impact the response to extracellular divalent concentration when creWT and creCasr^-/-^ neuron recordings were started at a membrane potential of −70 mV l (F (1, 29) = 0.2005, P = 0.6577). Post-hoc testing showed that excitability was increased in T_0.2_ compared with regardless which of the two holding currents were used (Figure 2D; Suppl. Table 7). The depolarization of Casr^-/-^ neurons in response to T_0.2_ was unexpected (Figure 1I) since initially CaSR-NALCN signaling was hypothesized as the mechanism for [Ca^2+^]_o_-dependent excitability(Lu et al., 2010). After injection of I_a_ to set the membrane potential to −70 mV, the switch from to T_0.2_ significantly depolarized the membrane potential (Figure 2E; Suppl. Table 8, Two-way RM ANOVA, F (1, 29) = 29.22, P < 0.0001) as did CaSR deletion (F (1, 29) = 4.874, P = 0.0353). Post-hoc testing indicate that the membrane potential in T_0.2_ was more depolarized in the creCasr^-/-^ than in creWT neurons (Figure 2B,E; - 65.6 ±1.6 mV vs −59.4 ± 2.4 mV, P = 0.0083). Taken together, these experiments indicate CaSR-NALCN signaling was not contributing to the difference in [Ca^2+^]_o_-dependent excitability between creWT and creCasr^-/-^ neurons but that these differences may be due to genotype-dependent differences in RMP or intrinsic excitability.

**Figure 2:**
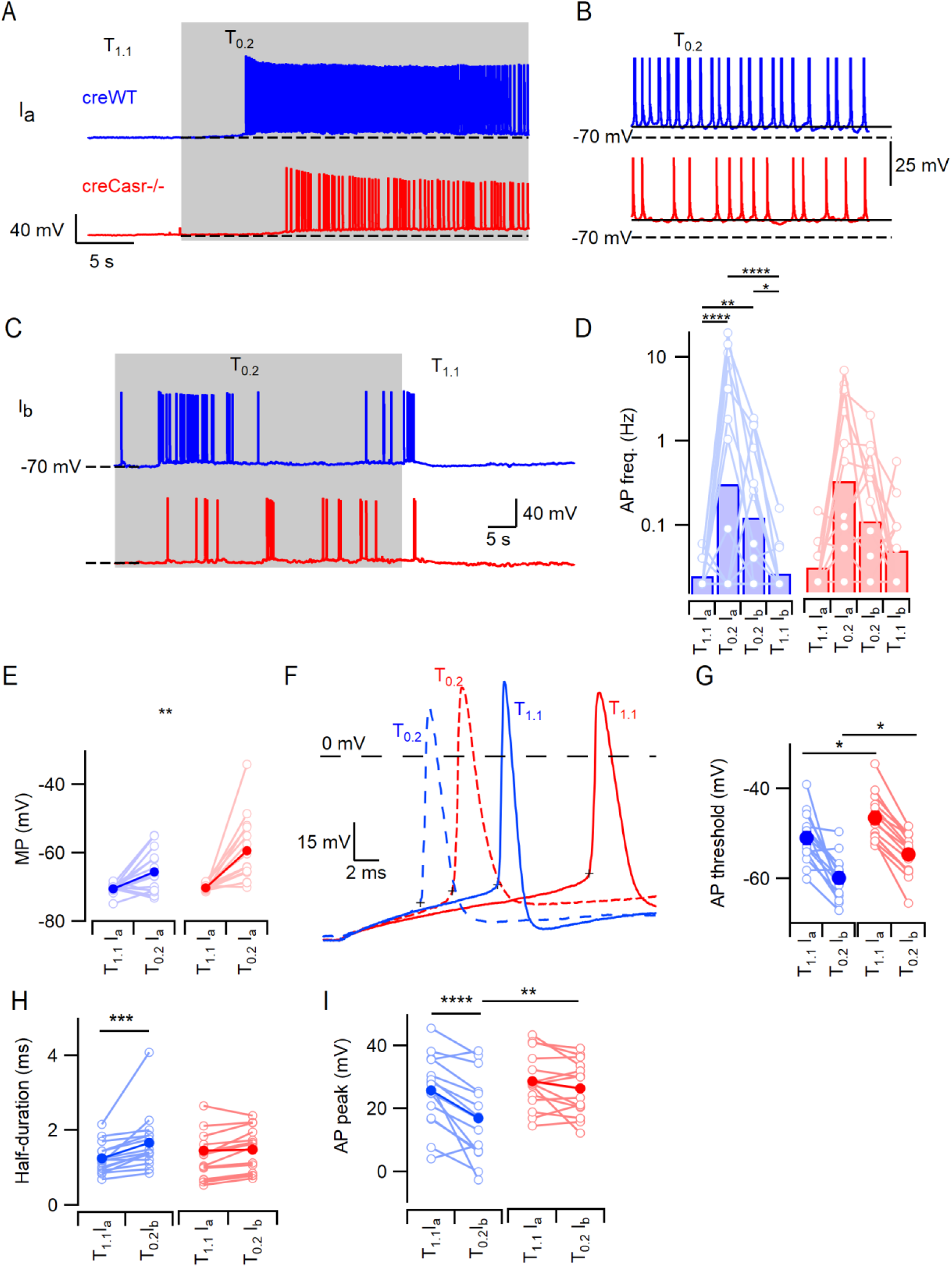
CaSR deletion does not affect [Ca^2+^]_o_-dependent excitability at equivalent membrane potential. A) Exemplary traces showing the [Ca^2+^]_o_-dependent increase in neuronal excitability following the switch from to T_0.2_ (change indicated by upper trace) in creWT (blue) and cre casr^-/-^ (red) neurons when initial membrane potentials matched at −70 mV. B) Expanded view of the final five seconds of traces in A illustrating sustained depolarization from −70 mV following T_0.2_ application. C) Exemplary traces showing the [Ca^2+^]_o_-dependent decrease in neuronal excitability following the switch from T_0.2_ to (change indicated by upper trace) in creWT (blue) and cre casr^-/-^ (red) neurons when initial membrane potentials matched at −70 mV. Same recording as A. D) Histogram of average [Ca^2+^]_o_-dependent changes in action potential frequency (Hz) in creWT (blue) and cre casr^-/-^ (red) neurons when initial voltage is −70 mV in (Ia) or T_0.2_ (Ib). Two-way RM ANOVA performed after logarithmic transformation indicates that reducing [Ca^2+^]_o_ increases the AP frequency (F (3, 87) = 17.97, P < 0.0001) similarly in creWT and creCasr^-/-^ neurons (F (1, 29) = _0.2_005, P = 0.6577). Post-hoc tests indicate significant differences between AP frequency in and T_0.2_ regardless of the holding current but not between AP frequency recorded at different holding currents and the same solutions (Ia or Ib; Supplementary Table?). E) Membrane potential depolarization following the switch to T_0.2_ from. Two-way RM ANOVA indicates that reducing [Ca^2+^]_o_ (F (1, 29) = 29.22, P < 0.0001) and CaSR deletion (F (1, 29) = 4.874, P = 0.0353) significantly depolarized the membrane potential but that there was no interaction (F (1, 29) = 4.055, P = 0.0534). Post-hoc testing indicate that membrane potentials were matched using current injection in (−70.5 ± 0.4 mV and −70.2 ± 0.2 mV for creWT and creCasr^-/-^ neurons respectively, P = 0.985) but different in T_0.2_ (−65.6 ± 1.6 mV and –59.4 ± 2.4 mV, P = 0.0083). F) Exemplar action potentials elicited by current injection from −70 mV in a creWT (blue) and a creCasr^-/-^ neuron (red) in solutions (unbroken) and T_0.2_ (broken). Action potential threshold is indicated by + for the first AP elicited by current injection (50 to 200 pA). G) Plot of average AP threshold in and T_0.2_ in creWT and creCasr^-/-^ neurons, elicited as per panel F here and in subsequent panels. Two-way RM ANOVA indicates that reducing [Ca^2+^]_o_ hyperpolarized the AP threshold (F (1, 25) = 51.66, P < 0.0001) whereas CaSR deletion had the opposite effect (F (1, 25) = 10.52, P = 0.0033). There was no interaction (Supplementary Table). Post-hoc tests indicate that the AP thresholds in solutions and T_0.2_ were depolarized similarly by CaSR deletion (5.3 ± 2.0 mV and 5.5 ± 2.0 mV, P = 0.020 and 0.017) in creWT and creCasr^-/-^ neurons respectively. H) Plot of average AP half-duration in and T_0.2_ in creWT and creCasr^-/-^ neurons. Two-way RM ANOVA indicates that reducing [Ca^2+^]_o_ prolonged the AP half-duration (F (1, 28) = 19.73, P = 0.0001). I) Plot of average AP peak in and T_0.2_ in creWT and creCasr^-/-^ neurons. The AP peaks were higher in T_1.1_ and in Casr-/- neurons (Supplementary Table 9C).

### Voltage-gated sodium channels contribute to [Ca^2+^]_o_-dependent excitability

Reversal of the [Ca^2+^]_o_-dependent depolarization did not completely block the increased excitability associated with the switch to T_0.2_ (Figures 1E,F & 2D) indicating another mechanism other than [Ca^2+^]_o_-dependent NALCN was responsible. We tested if voltage-gated channels were contributing by to [Ca^2+^]_o_-dependent excitability by examining action potential threshold in neurons held at a membrane potential of −70 mV. AP threshold was hyperpolarized by 8 mV on average following the change from to T_0.2_ in creWT and creCasr^-/-^ neurons (Figure 2F,G, Suppl. Table 9A; F (1, 25) = 51.66, P< 0.0001). Furthermore, the AP threshold was relatively depolarized in the creCasr^-/-^ neurons in and T_0.2_ (5.3 ± 2.0 mV (P = 0.020) and 5.5 ± 2.0 mV (P = 0.017) respectively), indicating creWT neurons possessed increased excitability and increased sensitivity to decreases in external divalent concentration (Figure 2F,G). The action potential half-width recorded under the same conditions, was also sensitive to deletion of CaSR and reduction of [Ca^2+^]_o_ (Figure 2H,I Suppl. Table 9B). ANOVA indicated that the switch to T_0.2_ from broadened AP half-width (F (1,28) = 19.7, P = 0.0001) but this effect was restricted to creWT (P = 0.0004) and not creCasr-/- (P = 0.117) neurons. The genotype and [Ca^2+^]_o_ interacted to both affect AP peak voltage (Figure 2I, Suppl. Table 9C; F (1, 28) = 6.76, P = 0.015) with the peak potential being reduced by T_0.2_ in the creWT (P < 0.0001) but not creCasr-/- neurons (P = 0.34).

We examined the properties of VGSCs and voltage-gated potassium channels (VGPCs) to determine the reason for the altered AP threshold. VGSCs were isolated in neocortical neurons and the current-voltage characteristics examined. Families of VGSC currents were activated in neurons after 2-4 weeks in culture. Maximum VGSC currents were elicited at −30 mV and averaged −8.0 ± 0.8 nA (n=7) and −8.8 ± 2.8 nA (n=6) in creWT and creCasr^-/-^ neurons respectively. The current-voltage curve shifted in a hyperpolarizing direction with the switch from to T_0.2_ but extensive neuronal processes limited the quality of the voltage-clamp and prevented useful analysis. We next examined VGSC gating in nucleated outside-out patches(Sather et al., 1992; Almog et al., 2018) to ensure better voltage control. VGSC currents were elicited by voltage steps from −80 mV (10 mV increments to 40 mV). In T_0.2_, the VGSC inactivation (see below) resulted in smaller currents that were more sensitive to depolarization (bold traces elicited by steps to −50 mV, Figure 3A) as previously observed (Frankenhaeuser and Hodgkin, 1957; Campbell and Hille, 1976; Armstrong and Cota, 1991). [Ca^2+^]_o_ sensitivity was confirmed in the normalized current-voltage plot for both creWT (blue, n= 8) and creCasr^-/-^ (red, n= 11) neurons (Figure 3A,C). VGSC current inactivation was studied using a test pulse to −20 mV, each of which was preceded by a conditioning step (100 ms) to between −140 mV and −20 mV. In we observed less inactivation than in T_0.2_ (Figure 3B, bold traces show currents elicited following prepulse to −80 mV). We compared the effects of [Ca^2+^]_o_ and CaSR deletion on VGSC current inactivation using plots of normalized conductance and measuring the half maximal voltage (V_0.5_) (circles, Figure 3D,E). The reduction in [Ca^2+^]_o_ (F (1, 18) = 56, P < 0.0001) left-shifted V_0.5_ (2-way RM ANOVA, Suppl. Table 10) but deletion did not CaSR (F (1, 18) = 0.563, P = 0.463). The switch from to T_0.2_ shifted V_0.5_ by −19 and −21 mV in creWT and creCasr^-/-^ respectively (P < 0.0001 for both). We also tested how the VGSC activation properties were affected by CaSR and [Ca^2+^]_o_. The peak inward VGSC currents (Figure 3B,D) were divided by the driving voltage and then plotted as conductance-voltage plots. The normalized conductance plots (squares, Figure 3E,F) indicate that the switch from to T_0.2_ significantly facilitated VGSC activation (V_0.5_ was hyperpolarized by 10 mV) (F (1, 17) = 98, P < 0.0001; 2-way RM ANOVA, Suppl. Table 11) consistent with VGSCs in other excitable cells (Hille, 2001). Switching from to T_0.2_ shifted V_0.5_ by −10 mV and −9 mV in creWT and creCasr^-/-^ neurons respectively (P < 0.0001 in both). Unexpectedly, the V_0.5_ for VGSC activation was also impacted by CaSR deletion, the relative depolarization reducing the likelihood of VGSC activation (F (1, 17) = 4.8, P = 0.04) in the Casr-/- neurons (Figure 3I) Overlap of the inactivation and activation conductance plots represents the voltage range over which persistent VGSC currents, or window currents, are likely to occur(Chadda et al., 2017). [Ca^2+^]_o_ reduction and CaSR hyperpolarized this region of overlap towards the RMP (Figure 3 F,G insets) increasing the likelihood that persistent VGSC currents were activated at resting membrane potential and therefore contributing to [Ca^2+^]_o_-dependent excitability.

**Figure 3.**
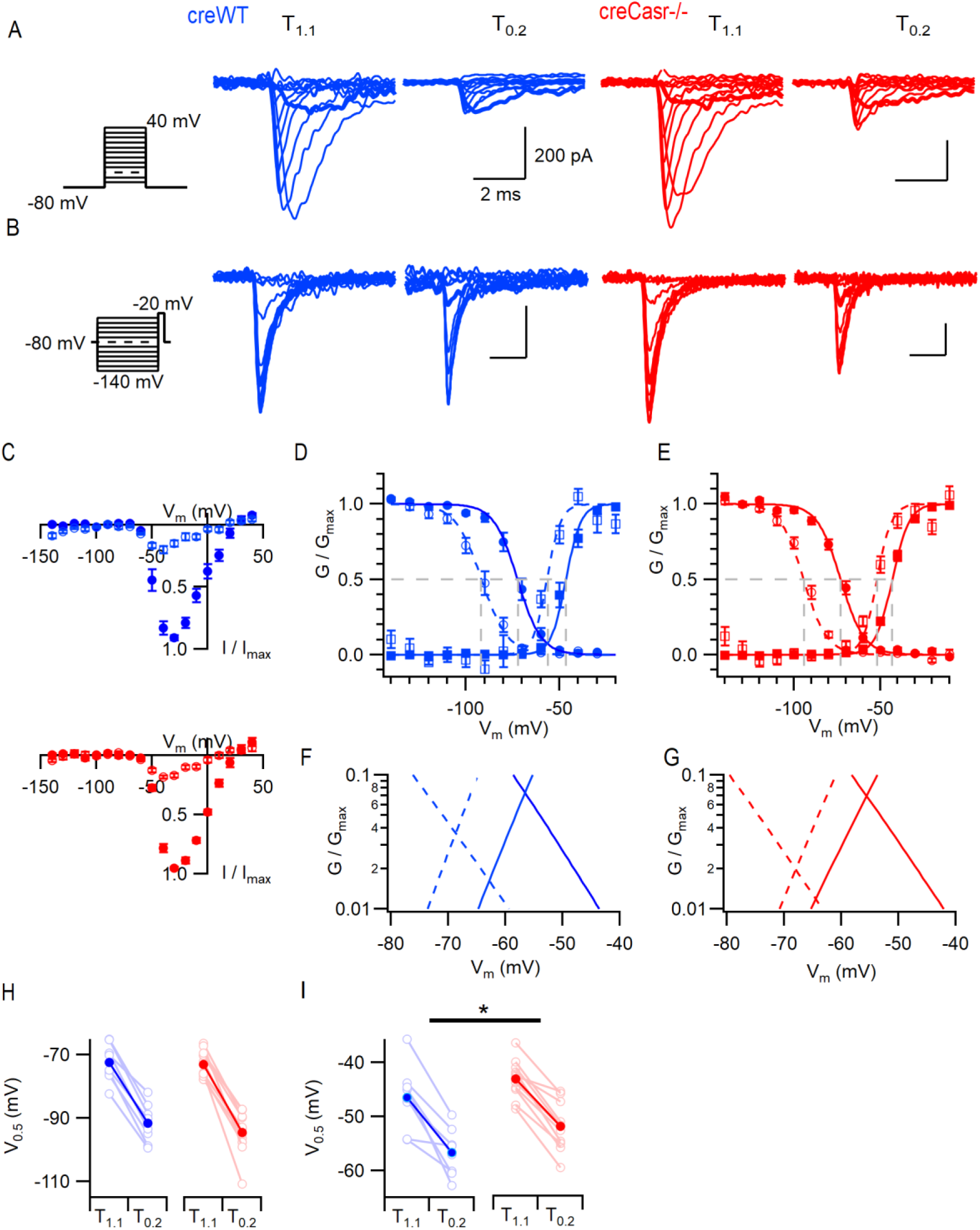
CaSR deletion and [Ca^2+^]_o_ affect VGSC current gating. A) Exemplary traces showing VGSC currents activated by voltage steps from −80 in 10 mV increments (left), in nucleated patches isolated from creWT (blue) and creCasr-/- (red) neurons in solutions and T_0.2_. The VGSC currents elicited by 10 ms depolarizations to - 50 mV (bold) were greater following the switch to T_0.2_. B) Exemplary traces showing VGSC currents activated by voltage steps to −20 mV following a 100 ms conditioning step (left), in the same patches as A) using solutions and T_0.2_. The VGSC currents elicited following conditioning steps to −80 mV (bold) were smaller following the switch to T_0.2_. C) Current-voltage plots of average normalized VGSC currents in nucleated patches from creWT (n = 8) and creCasr-/- (n = 11) neurons in (filled circles) and T_0.2_ (open circles). Currents were normalized using the maximum VGSC current in each recording. D) Plot of average normalized conductance versus voltage in patches from creWT neurons for activation (square, n = 8) and inactivation (circle, n = 8) in solutions (filled) and T_0.2_ (open). Boltzmannn curves are drawn using average values from individual fits and gray broken lines indicate V_0.5_ values for each condition. E) Plot of average normalized conductance versus voltage in patches from creCasr^-/-^ neurons for activation (square, n = 11) and inactivation (circle, n = 12) in solutions (filled) and T_0.2_ (open). Boltzmannn curves are drawn using average values from individual fits and gray broken lines indicate V_0.5_ values for each condition. Inset shows plot expanded to emphasize voltage dependence of the window currents. F and G represent the plots of D and E expanded to emphasize the voltage dependence of the window currents. H) Histogram showing V_0.5_ for VGSC inactivation in and T_0.2_ in patches from creWT and creCasr^-/-^ neurons. I) Histogram showing V_0.5_ for VGSC activation in and T_0.2_ in patches from creWT and creCasr^-/-^ neurons.

VGPC currents were isolated and recorded in creWT and creCasr^-/-^ neurons in and T_0.2_ solutions after blocking contaminating currents. Currents were elicited by a series of 60 ms steps from −70 mV to 60 mV in 10 mV increments (Figure 4). The VGPC current amplitudes were measured at the peak and at the end of the depolarizing step (normalized to the value at 60 mV in). Neither the peak nor end current were affected by reduction of the external divalent concentration or by deletion of CaSR (Figure 4) over the range of voltages. The currents activated at 60 mV were similarly unaffected (Figure 4, Two-way RM ANOVA ((3, 57) = 1.347, P= 0.2683 and (1, 19) = 1.231, P= 0.2811, Suppl. Table 12). These data indicate that VGPCs are not involved in [Ca^2+^]_o_-dependent excitability in neocortical neurons.

**Figure 4.**
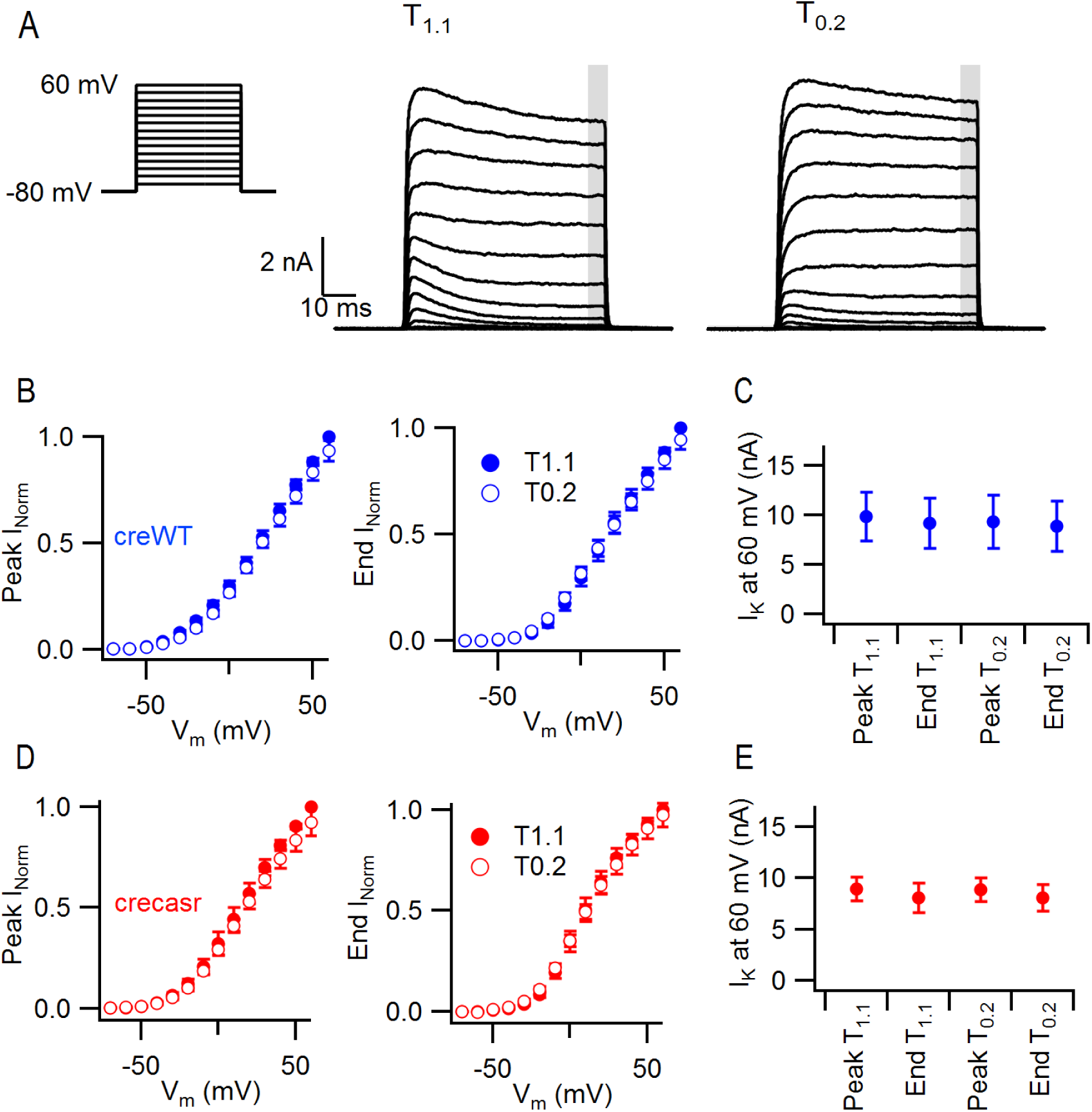
CaSR deletion and [Ca^2+^]_o_ do not significantly affect VGPC current gating. A) Exemplary traces showing VGPC currents activated by voltage steps from −80 in 10 mV increments (left), in a creWT neuron in solutions and T_0.2_. The outward currents elicited by the 50 ms voltage step were measured at peak and at the end of the step (average of last 5 ms indicated by gray bar). B) Current voltage-plot of average normalized VGPC currents (n = 10) in creWT neurons in (filled circles) and T_0.2_ (open circles) at peak or end of step. Currents were normalized using the maximum outward current in each condition here and below. C) Peak and end outward currents at 60 mV elicited in same neurons as B. Two-way RM ANOVA indicates that peak and outward currents were not different in or T_0.2_ ((3, 57) = 1.347, P= 0.2683 nor were they affected by CaSR deletion (data from E, (1, 19) = 1.231, P= 0.2811). D) Current voltage-plot of average normalized VGPC currents (n = 11) in creCasr-/- neurons in (filled circles) and T_0.2_ (open circles) at peak or end of step. E) Peak and end outward currents at 60 mV elicited in same neurons as D.

### VGSCs are the dominant contributor to [Ca^2+^]_o_-dependent currents

To compare the contributions of VGSCs and NALCN to the [Ca^2+^]_o_-dependent depolarization seen in neocortical neurons (Figure 2) we measured the size of the currents elicited at −70 mV in neurons following the switch from to T_0.2_. We used conWT neurons to avoid potential confounding Cre-dependent effects (Qiu et al., 2011). Since NALCN is resistant to the VGSC blocker tetrodotoxin (TTX) (Lu et al., 2007; Swayne et al., 2009) but Gd^3+^ (10 µM) inhibits NALCN and VGSCs (Elinder and Arhem, 1994; Li and Baumgarten, 2001; Lu et al., 2009), we were able to pharmacologically separate the contributions of VGSCs and NALCN to the basal current following the switch from to T_0.2_ (−31 ± 3 pA, n=13; Figure 5A-C). Addition of a saturating concentration of TTX (1 µM) in T_0.2_ inhibited a persistent inward current within a few seconds in all but one of the recordings (Figure 5A-C), consistent with VGSCs contributing to the inward current elicited by T_0.2_. Switching to plus TTX produced minimal change in the basal current on average (Figure 5C). However, in some neurons elicited an outward current (Figure 5A,C) whereas in others there was an inward current (Figure 5B,C) indicating the presence of two types of TTX resistant [Ca^2+^]_o_-dependent pathways. Presumably NALCN was contributing to the [Ca^2+^]_o_-dependent TTX-resistant effect observed in Figure 5A. Co-application of Gd^3+^ (10 µM) following block of VGSCs with TTX, resulted in a small inward deflection of the average basal current in solution and largely inhibited sensitivity to concomitant decreases in [Ca^2+^]_o_ (Figure 5A-C). The reduced sensitivity of neurons to the reduction of [Ca^2+^]_o_ in the presence of TTX, suggests that VGSCs are a major contributor to the depolarizing current elicited by low [Ca^2+^]_o_.

**Figure 5.**
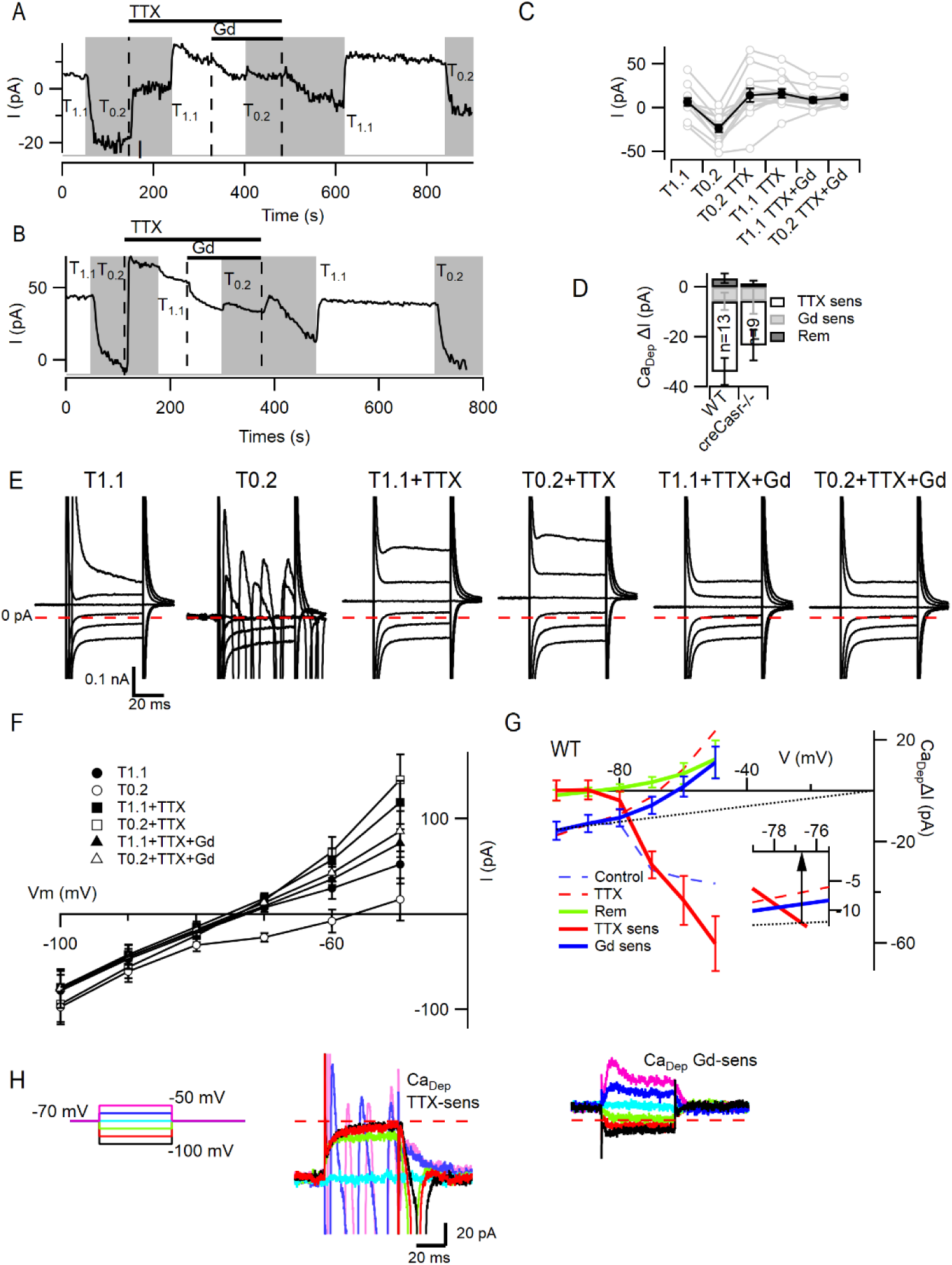
VGSC current activation by decreased [Ca^2+^]_o_. A,B) Plots illustrating the responses of the basal currents in two WT neurons during application of and T_0.2_ before and during TTX or TTX and Gd^3+^. Average basal currents were measured over 50 ms every 2 s with and T_0.2_ application indicated by vertical shading (gray represents T_0.2_) and blockers application by horizontal bars and broken vertical lines. C) Plot of average basal current measurements (filled circles) and individual neurons (open circles) in each solution condition in conWT (n=13) neurons. Each basal current represents the average value recorded during last 20 s of the specific solution application. D) Average [Ca^2+^]_o_ dependent basal currents sensitive to TTX and Gd^3+^ calculated by subtraction of data in C and the remaining [Ca^2+^]_o_ dependent current after application of both blockers for conWT neurons. E) Exemplar traces of currents elicited by 50 ms voltage steps between −100 and −50 mV during application of solutions described in C. F) Plots of the average currents over the last 5 ms of each voltage step in all six solutions for conWT (n = 13). G) Plots of the average [Ca^2+^]_o_ dependent currents derived by subtraction of WT data (F) resolved as total or control (broken blue), in the presence of TTX (broken red), and in the presence of TTX and Gd^3+^ (remainder green). The TTX-sensitive (solid red), Gd^3+^-sensitive (solid blue) and NALCN (dotted line) component currents were obtained by further subtraction. Inset shows expanded view at intercept of TTX-sensitive and NALCN components. H) Exemplars of the TTX- and Gd^3+^-sensitive [Ca^2+^]_o_ dependent currents. Broken red line represents zero current line.

Using serial subtraction of the basal currents (Figure 5C), we compared the size of the TTX-sensitive (−28.2 ± 5.3 pA), Gd^3+^-sensitive (−5.7 ± 3.4 pA) and remaining (3.4 ± 2.0 pA) [Ca^2+^]_o_-dependent currents (Figure 5D; RM-ANOVA, F (1.495, 17.94) = 13.30, P = 0.0007, Suppl. Table 13). Multiple comparison testing indicated that the TTX-sensitive [Ca^2+^]_o_-dependent current was greater than the Gd^3+^-sensitive (P = 0.039) and remaining [Ca^2+^]_o_-dependent currents (P = 0.0009; Suppl. Table 14). Similar differences in the relative sizes of the TTX-, Gd^3+^-, and remainder [Ca^2+^]_o_-dependent basal current currents were also observed in Casr^-/-^neurons (Figure 5D). While there were rare neurons in which there was a larger Gd^3+^-sensitive current (Figure 5C) the reduced sensitivity of neurons to the reduction of [Ca^2+^]_o_ in the presence of TTX, confirms that VGSCs are the major contributor to the depolarizing current elicited by low [Ca^2+^]_o._.

In a fraction of the neurons, an inward deflection of the basal current occurred when [Ca^2+^]_o_ was increased in the presence of TTX (Figure 5B,C) which contrasted with the outward current expected from NALCN deactivation (Figure 5A). We examined the voltage-dependence of the contributions of VGSCs, NALCN, and this second [Ca^2+^]_o_-dependent TTX-resistant current to better characterize [Ca^2+^]_o_-dependent excitability. We used 50 ms voltage steps between −100 and −50 mV and averaged the current over the last 5 ms of the step. Three additional major effects are illustrated by the exemplar current traces (Figure 5E). First, in the absence of blockers, the switch from to T_0.2_ substantially increased the number of large, rapidly inactivating inward currents even at −70 mV following hyperpolarizing steps. Second, in TTX, low [Ca^2+^]_o_ increased the linear inward and rectifying outward currents. Third, in the presence of TTX and Gd^3+^ changing between and T_0.2_ had little effect suggesting Gd^3+^ is blocking both NALCN and the second [Ca^2+^]_o_-dependent TTX-resistant current. These observations were confirmed in the average current-voltage plots (Figure 5F) where it is clear that at −80 to −100 mV the major [Ca^2+^]_o_-dependent currents are inward and resistant to TTX and sensitive to Gd^3+^ whereas at −70 to −50 mV the largest [Ca^2+^]_o_-dependent currents are TTX-sensitive. The [Ca^2+^]_o_-dependent effects were calculated by subtracting the currents recorded in from those in T_0.2_ under control conditions (Figure 5G, broken red), in the presence of TTX (broken blue) and TTX plus Gd^3+^ (solid green). The TTX-sensitive (solid red) and Gd^3+^-sensitive (solid blue) [Ca^2+^]_o_-dependent currents were obtained by additional subtraction (broken red minus broken blue and broken blue minus green). The average [Ca^2+^]_o_-dependent current carried by VGSCs only became evident once the neurons were depolarized above −80 mV (Figure 5G). The time course of deactivation of the persistent [Ca^2+^]_o_-dependent VGSC currents was observed following hyperpolarization from −70 mV (Figure 5H, middle). At more negative potentials, the Gd^3+^-sensitive current accounted for all of the [Ca^2+^]_o_-dependent current and traces showed an ohmic voltage dependence (Figure 5G). However, the Gd^3+^-sensitive current reversed at −60 mV and outward currents were elicited by steps to −60 and −50 mV that exhibited a voltage-dependent activation and inactivation (Figure 5H, right-hand). This is consistent with the Gd^3+^-sensitive current consisting of the sum of NALCN and an outward voltage-dependent current. Assuming conservatively that all of the Gd^3+^-sensitive current at −100 mV could be attributed to NALCN and employing the channel’s linear voltage-dependence and zero mV reversal potential(Lu et al., 2007; Lu et al., 2010), then the amplitude of NALCN currents could be estimated over the voltage range −100 to 0 mV (broken black line, Figure 5G). By interpolation (Figure 5G, inset), the contribution of NALCN and VGSCs to [Ca^2+^]_o_-dependent currents were equal at ∼-77 mV with the contribution from VGSCs increasing with depolarization. A similar analysis of [Ca^2+^]_o_-dependent currents in creCasr^-/-^ neurons indicated that the contribution of VGSCs was greater than that of NALCN once membrane potentials were depolarized beyond - 72 mV (Figure 5 Supplement). These data indicate that [Ca^2+^]_o_-dependent currents around the resting membrane that contribute to [Ca^2+^]_o_-dependent excitability are mainly attributable to VGSCs.

### VGSCs are the dominant contributor to [Ca^2+^]_o_-dependent excitability

The complex architecture of neocortical neurons restricted our ability to clamp the membrane potential following the activation of large, rapid VGSC currents. Thus we re-examined the contribution of VGSCs and NALCN to the depolarizations that mediate [Ca^2+^]_o_-dependent excitability in current clamp recordings from conWT neurons. Consistent with earlier experiments (Figure 2), switching from to T_0.2_ depolarized the membrane potential from −70 mV by 7.2 ± 1.5 mV (n = 12) and increased spontaneous action potential firing in pharmacologically isolated neurons (Figure 6A,B). We used TTX and Gd^3+^ to measure the contributions of VGSCs and NALCN respectively to these [Ca^2+^]_o_-dependent depolarizations. TTX blocked action potential generation, as expected, but also hyperpolarized the membrane potential indicating that VGSCs were open in T_0.2_ (Figure 6A1) and (Figure 6A2). The switch from T_0.2_ to in TTX resulted in a hyperpolarization, consistent with [Ca^2+^]_o_-dependent NALCN closure, in some neurons (Figure 6A1 lower trace and B). Other neurons depolarized with the switch to (Figure 6A2 lower trace and B) consistent with a [Ca^2+^]_o_-dependent outward current similar to that observed in Figure 5B,C. On average the [Ca^2+^]_o_-dependent depolarization was almost entirely prevented by TTX or TTX and Gd^3+^ co-application (Figure 6B). The [Ca^2+^]_o_-dependent depolarizations in conWT neurons were different for blocker type (1-way RM ANOVA, F (1.219, 13.41) = 12.83, P = 0.0022, Suppl. Table 15) and more sensitive to TTX than Gd^3+^ or following co-application (P = 0.022 and 0.0028 respectively, Supp. Table 16). On average VGSCs accounted for 93% of the depolarization that triggers [Ca^2+^]_o_-dependent excitability in WT neurons starting at −70 mV (Figure 6C) and we observed a similar pattern in Casr^-/-^ neurons (Figure 6C).

**Figure 6.**
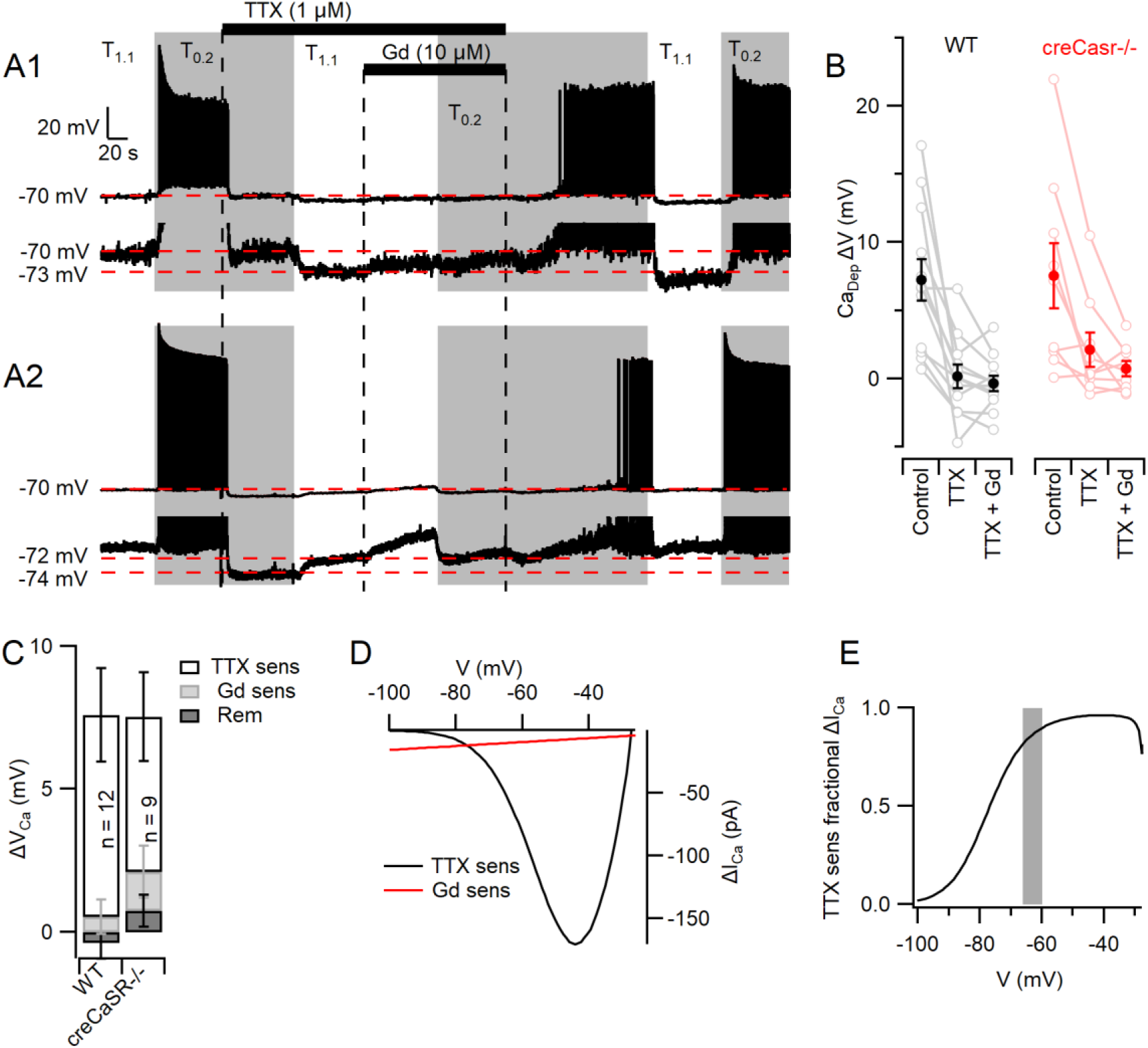
[Ca^2+^]_o_-dependent depolarization is almost entirely mediated via VGSCs. A) The response of the membrane potential in two WT neurons during application of and T_0.2_ before and during TTX or TTX and Gd^3+^. and T_0.2_ application is indicated by vertical shading (gray represents T_0.2_) and blocker applications by horizontal bars and broken vertical lines. The broken red line indicates −70 mV. Voltage-expanded view of the trace illustrates that in the presence of TTX, hyperpolarization (A1) and depolarization (A2) may occur following the switch to. Membrane potential values highlighted by broken red lines. B) Plot of average (filled circles) and individual (open circles) Ca^2+^-dependent voltage changes (filled circles) following the switch from to T_0.2_ (by subtraction of average between-spike membrane potential over the last 10 sec of each solution application). Each solution applied to conWT (n=12) and creCasr-/- (n = 9) neurons. C) Average [Ca^2+^]_o_ dependent voltage changes sensitive to TTX and Gd^3+^ calculated by subtraction of data in B, and the remaining [Ca^2+^]_o_ dependent voltage after application of both blockers. D) Estimates of the average relative size of the [Ca^2+^]_o_ dependent NALCN and VGSC currents in neocortical neurons between −100 and −30 mV. NALCN values from Fig 5G. The [Ca^2+^]_o_-dependent VGSC currents were estimated as follows: the products of the VGSC activation and inactivation conductance plots were calculated for T_0.2_ using the average Boltzmann curves in Figure 3. These were converted to currents (I= driving voltage x conductance), and scaled to match the average [Ca^2+^]_o_ dependent TTX-sensitive current at −70 mV. The current generated in T_0.2_ minus that generated in ΔI_Ca_) was plotted against membrane voltage. E) Plot of the fraction of the average [Ca^2+^]_o_ dependent depolarizing current carried by VGSC derived from D. The change in average resting membrane potential recorded in Figure 1 is indicated by the gray bar.

Next we estimated the average relative contributions of the [Ca^2+^]_o_-dependent NALCN and VGSC currents over a wider physiological voltage range. [Ca^2+^]_o_-dependent NALCN currents were extrapolated from −100 mV, where contaminating currents appear minimal (Figure 5G) and compared with the [Ca^2+^]_o_-dependent VGSC currents predicted from scaled conductance plots (Figure 3D). The VGSC currents were the major contributor to [Ca^2+^]_o_-dependent currents over the −77 to −30 mV voltage range (Figure 6 D,E) suggesting VGSCs are likely to dominate those of NALCN at resting membrane potentials observed in T_1.1_ and T_0.2_ (Figure 6E, gray bar).

## Discussion

Extracellular calcium concentration regulates both synaptic transmission and intrinsic neuronal excitability, thereby strongly affecting the probability of action potential generation. Consequently, physiological and pathological changes in [Ca^2+^]_o_ will impact neuronal computation in a complex manner. We have investigated the mechanisms underlying [Ca^2+^]_o_-dependent changes in intrinsic neuronal excitability and tested if CaSR is transducing decreases in [Ca^2+^]_o_ into NALCN-mediated depolarizations to trigger action potentials (Lu et al., 2010; Philippart and Khaliq, 2018). We found no evidence that this specific mechanism was active in neocortical neurons (Figure 2). Instead, we determined that the vast majority of [Ca^2+^]_o_ dependent neuronal excitability was mediated via VGSCs in three ways. First, decreased [Ca^2+^]_o_ activated VGSCs at the resting membrane potential and depolarized the membrane towards the action potential threshold (Figure 6). Second, decreased [Ca^2+^]_o_ hyperpolarized the expected VGSC window current towards the membrane potential increasing the likelihood of action potential generation (Figure 3). Third, deletion of CaSR modulated VGSC gating by synergistically increasing the sensitivity of the currents to depolarization via an unidentified mechanism (Figure 3). CaSR also indirectly affected action potential generation by modestly depolarizing the membrane potential (Figure 1). While the actions of [Ca^2+^]_o_ on VGSCs were responsible for the vast majority of the [Ca^2+^]_o_ dependent neuronal excitability, using Gd^3+^ we isolated small [Ca^2+^]_o-_dependent inward currents in about half of the neurons (Figure 5). These small Gd^2+^-sensitive currents presumably reflected activation of NALCN, and were unaffected by CaSR deletion, but their relatively small size compared to TTX-sensitive [Ca^2+^]_o-_dependent inward currents indicate that they would be minor contributors to [Ca^2+^]_o_-dependent excitability compared to VGSCs (Figures 5,6).

The fractions of the [Ca^2+^]_o_-dependent currents and depolarizations that were sensitive to TTX were surprisingly large compared to those that were Gd^3+^-sensitive (Figures 5D,6C) indicating the relative importance of VGSC- and NALCN-mediated contributions to [Ca^2+^]_o_-dependent excitability respectively. The resistance of NALCN to TTX (Lu et al., 2007; Swayne et al., 2009) reassures that the relatively large TTX-sensitive component is due to selective block of VGSC. Persistent subthreshold VGSC currents have been shown to determine spiking rates in other central neurons (Taddese and Bean, 2002; Gorelova and Seamans, 2015) and so the increased VGSC currents we observed in T_0.2_ are well-positioned to explain the increased action potential frequency (Figure 6). We are unable to determine from these experiments which neuronal compartment is most affected by the change in [Ca^2+^]_o_(Gorelova and Seamans, 2015). However, the physiological impact of VGSC-mediated [Ca^2+^]_o-_dependent excitability will be enormous overall because of the dynamic nature of [Ca^2+^]_o_ in vivo where it decreases from basal (1.1-1.2 mM) by 30-80%(Nicholson et al., 1978; Ohta et al., 1997; Pietrobon and Moskowitz, 2014). The overall computational effects of physiological decrements in [Ca^2+^]_o_ will be complex because the increased action potential generation due to changes on VGSCs (Figure 3,6) will be confounded by the impact of reduced Ca^2+^ entry through VACCs (Hess et al., 1986; Weber et al., 2010), reduced synaptic transmission (Neher and Sakaba, 2008), and altered CaSR-mediated signaling at the nerve terminal (Phillips et al., 2008; Vyleta and Smith, 2011).

A number of questions arise from our studies. Why was NALCN essential to effect [Ca^2+^]_o_-dependent excitability in hippocampal(Lu et al., 2010) but not neocortical neurons (Figure 2)? Since action potentials were elicited by divalent concentration changes alone in neocortical neurons (Figure 2) whereas the response to current injections was the measure of excitability in hippocampal neurons(Lu et al., 2010) could the differences reflect different experimental protocols. We cannot exclude this possibility however, it is unclear how NALCN facilitated acute [Ca^2+^]_o_-dependent excitability during depolarizing injections from matched membrane potentials since the current injections were expected to produce similar or smaller depolarizations given NALCN is relatively voltage insensitive (Lu et al., 2009). Could the increased action potential number seen in WT compared to NALCN deficient neurons(Lu et al., 2010) reflect other indirect actions of a persistent leak current (Sokolov et al., 2007) or compensatory changes in ion channel activity(Jun et al., 1999)? It is also surprising that deletion of NALCN or UNC-79 completely ablated [Ca^2+^]_o_-dependent excitability in hippocampal neurons (Lu et al., 2010) since, like other excitable cells, hippocampal VGSC currents are sensitive to changes in [Ca^2+^]_o_ (Isaev et al., 2012). Consequently, if the UNC79-UNC80-NALCN pathway modulates VGSC function this could explain how loss of NALCN or UNC-79 could delete acute [Ca^2+^]_o_-dependent changes in VGSC function and excitability. NALCN appears to transduce [Ca^2+^]_o_- and G-protein dependent excitability in other neurons(Philippart and Khaliq, 2018) but GPCRs other than CaSR may be involved(Kubo et al., 1998; Tabata and Kano, 2004). Further characterization of the UNC79-UNC80-NALCN signaling pathway is essential given the major changes in neurological function that have been described following mutations of NALCN or upstream co-molecules such as UNC79 and UNC80(Stray-Pedersen et al., 2016; Bourque et al., 2018; Kuptanon et al., 2019).

In a small fraction of the neocortical neurons (Figures 5,6) there was a modest inward current or depolarization with the application of low calcium once VGSCs had been blocked. In a few cases they were sensitive to 10 µM Gd^3+^ consistent with a NALCN-mediated effect and those that were resistant were consistent with other [Ca^2+^]_o_-dependent non-selective cation channels(Ma et al., 2012). Deletion of CaSR, increased [Ca^2+^]_o_-dependent depolarizations (Figure 2E, 6B) indicating [Ca^2+^]_o_-dependent non-selective cation channels are perhaps upregulated in the absence of CaSR. While CaSR-NALCN signaling did not contribute to [Ca^2+^]_o_-dependent excitability in neocortical neurons (Figures 2 and 6) it was clear that Casr^-/-^ neurons were substantially less sensitive to changes [Ca^2+^]_o_ (Figure 1A,B). The reduced [Ca^2+^]_o_ sensitivity in these neurons is attributable to the combination of altered VGSC gating (Figure 3E,F) and the hyperpolarized RMP (Figure 1I). Although CaSR did not affect the amplitude of the shift in V_0.5_ following the switch to T_0.2_, V_0.5_ in was hyperpolarized by loss of CaSR (Figure 3EF). Could CaSR stimulation activate G-proteins and regulate the basal V_0.5_ for VGSC currents (Figure 3E,F)? In neocortical neurons, G-protein activation hyperpolarized VGSC gating and this was blocked by GDPβS (Mattheisen et al., 2018) which is inconsistent with the effect we observed here. Other possible explanations are that CaSR regulates VGSC subunit expression or post translational modification (Cantrell et al., 1996; Zhang et al., 2019). Loss of CaSR could hyperpolarize the neocortical neurons (Figure 1I) by increasing the function of depolarizing components or inhibiting hyperpolarizing elements, but it is unclear which of the many ion channels or pumps mediate this effect on RMP(Tavalin et al., 1997; Talley et al., 2001; Bean, 2007; Harnett et al., 2015; Hu and Bean, 2018).

Overall our studies indicate that [Ca^2+^]_o_-dependent excitability in neurons is largely attributable to actions of extracellular calcium on the VGSC function. Given the dynamic nature of brain extracellular calcium, this mechanism is likely to impact neuronal signaling greatly under physiological and pathological conditions. CaSR-dependent reduction of VGSC sensitivity to membrane potential adds further complexity to extracellular calcium signaling and identifies another potential mechanism by which CaSR stimulation increases neuronal death following stroke and traumatic brain injury(Kim et al., 2013; Hannan et al., 2018).

## Materials and Methods

### Genotyping and CaSR mutant mice

ConWT animals were obtained from a 20 year old colony consisting of a stable strain of C57BL/6J and 129×1 mice. The creCasr-/- mice were generated by breeding floxed Casr (Chang et al., 2008) and nestin Cre mice (B6.Cg-Tg (Nes-cre)1Kln/J, The Jackson Laboratory) as described previously (Sun et al., 2018). The lox sites were positioned to delete Casr exon 7 which resulted in the loss of Casr expression(Chang et al., 2008) and the nestin promoter was designed to ensure floxing occurred in neuronal and glial precursors. The creWT mice were generated by crossing mice that did not contain the flox Casr mutation but did express the nestin Cre mutation. The creCasr-/- and creWT mice were all generated using a background C57BL/6J and 129S4 strain. Tail DNA extraction was performed using the Hot Shot Technique with a 1-2 hour boil(Montero-Pau et al., 2008). The presence or absence of the flox Casr mutation and Cre mutation were confirmed by PCR for each mouse. MoPrimers used for cre PCR were: Nes-Cre 1: GCAAAACAGGCTCTAGCGTTCG, Nes-Cre 2: CTGTTTCACTATCCAGGTTACGG; run on a 1% agarose gel. Primers for lox PCR were: P3U: TGTGACGGAAAACATACTGC, Lox R: GCGTTTTTAGAGGGAAGCAG; run on a 1.5% agarose gel (Chang et al., 2008). Successful deletion of Casr in the neocortical cultures was confirmed by measuring mRNA expression levels with the QuantStudio12K Flex Real-time PCR System (Applied Biosystems)and the TaqMan mouse probe set to Casr (Mm00443377_m1) with ActB (Mm04394036_g1) as the endogenous control (Supplementary Figure 1).

### Neuronal Cell Culture

Neocortical neurons were isolated from postnatal day 1–2 mouse pups of either sex as described previously(Phillips et al., 2008). All animal procedures were approved by V.A. Portland Health Care System Institutional Animal Care and Use Committee in accordance with the U.S. Public Health Service Policy on Humane Care and Use of Laboratory Animals and the National Institutes of Health Guide for the Care and Use of Laboratory Animals. Animals were decapitated following induction of general anesthesia with isoflurane and then the cerebral cortices were removed. Cortices were incubated in trypsin and DNase and then dissociated with a heat polished pipette. Dissociated cells were cultured in MEM plus 5% FBS on glass coverslips. Cytosine arabinoside (4 µM) was added 48 hours after plating to limit glial division. Cells were used, unless otherwise stated after ≥14days in culture.

### Electrophysiological Recordings

Cells were visualized with a Zeiss IM 35 inverted microscope. Whole-cell voltage-and current-clamp recordings were made from cultured neocortical neurons using a HEKA EPC10 USB amplifier. Except where stated in the text, extracellular Tyrodes solution contained (mM) 150 NaCl, 4 KCl, 10 HEPES, 10 glucose, 1.1 MgCl_2_, 1.1 CaCl_2_, pH 7.35 with NaOH. Calcium and magnesium were modified as described in the Figure legends Synaptic transmission was blocked by the addition of (in µM) 10 CNQX, 10 Gabazine, and 50 APV to the bath solution. Most recordings were made using a potassium gluconate intracellular solution containing (mM) 135 K-gluconate, 10 HEPES, 4 MgCl_2_, 4 NaATP, 0.3 NaGTP, 10 phosphocreatine disodium, pH 7.2 with KOH hydroxide. In nucleated patch experiments the pipette solution contained (in mM) 113 Cesium methane sulfonate, 1.8 EGTA, 10 HEPES, 4 MgCl_2_, 0.2 CaCl_2,_ 4 NaATP, 0.3 NaGTP, 14 phosphocreatine disodium, pH 7.2 with TEA hydroxide. Electrodes used for recording had resistances ranging from 2 to 7 MΩ. Voltages have been corrected for liquid junction potentials. All experiments were performed at room temperature (21-23 °C).

### Data Acquisition and Analysis

Whole-cell voltage-and current-clamp recordings were made using a HEKA EPC10 USB amplifier, filtered at 2.9 kHz using a Bessel filter, and sampled at 20 kHz during injection protocols and 10 kHz during continuous acquisition. Analysis was performed using Igor Pro (Wavemetrics, Lake Oswego, OR) and Minianalysis (Synaptosoft). Data values are reported as mean ± SEM. Statistical tests were performed using GraphPad Prism (6) and P-values < 0.05, 0.01, 0.001, and 0.0001 were indicate with *, **, ***, and ****. All post-hoc tests were Sidak compensated for multiple comparisons. Data were log-transformed to improve normalization in Figure 2D. To ensure non-zero values, minimize bias, and allow logarithmic transformation, each AP frequency measurement was increased by 0.02 as the duration of the recording at −70 mV was 50 s.

### Solution Application

Solutions were applied by gravity from a glass capillary (1.2 mm outer diameter) placed ∼1 mm from the neuron under study. Solutions were switched manually using a low dead volume manifold upstream of the glass capillary. CNQX and Gabazine were supplied by Abcam. KB-R7943 Mesylate was supplied by Tocris. Creatine Phosphate was supplied by Santa Cruz Biotech. Cinacalcet was supplied by Toronto Research Chemicals and Tetrodotoxin by Alomone Other reagents were obtained from Sigma-Aldrich.

## Acknowledgements

This work was supported by grants awarded by U.S. Department of Veterans Affairs (BX002547) and NIGMS (GM134110) to SMS. We thank Dr Wenyan Chen for performing the experiments on potassium channel currents and Dr Glynis Mattheisen, Dr Brian Jones, Ms Natasha Baas-Thomas, and Ms Maya Feldthouse for helpful discussion and comments on the manuscript. Thanks to Drs Chris Harrington and Brittany Daughtry of the OHSU Gene Profiling Shared Resource which performed RNA isolation, quality assessments, and qPCR assays. The authors declare no competing financial interests. The contents do not represent the views of the U.S. Department of Veterans Affairs or the United States Government.

---

**Supplementary Figure 1.**
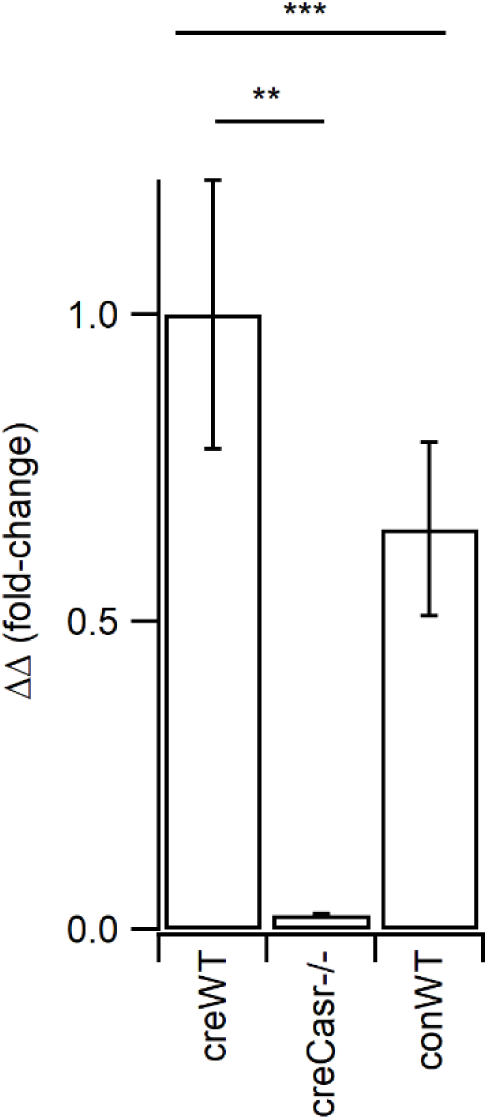
Casr expression levels shown as delta delta with actin used as normalizing gene. Average values of 1.0, 0.02, and 0.65 for creWT, creCasr-/-, and conWT respectively with each genotype reflecting the average data from six cultures each represented by triplicate samples. The Kruskal-Wallis test indicates differences between genotypes (P = 0.0002) with Dunn’s multiple comparison showing Casr expression levels are lower in creCasr-/- (P = 0.0016), but not conWT (P = 0.7739), than in creWT neocortical cultures.

**Figure 3 Supplementary figure.**
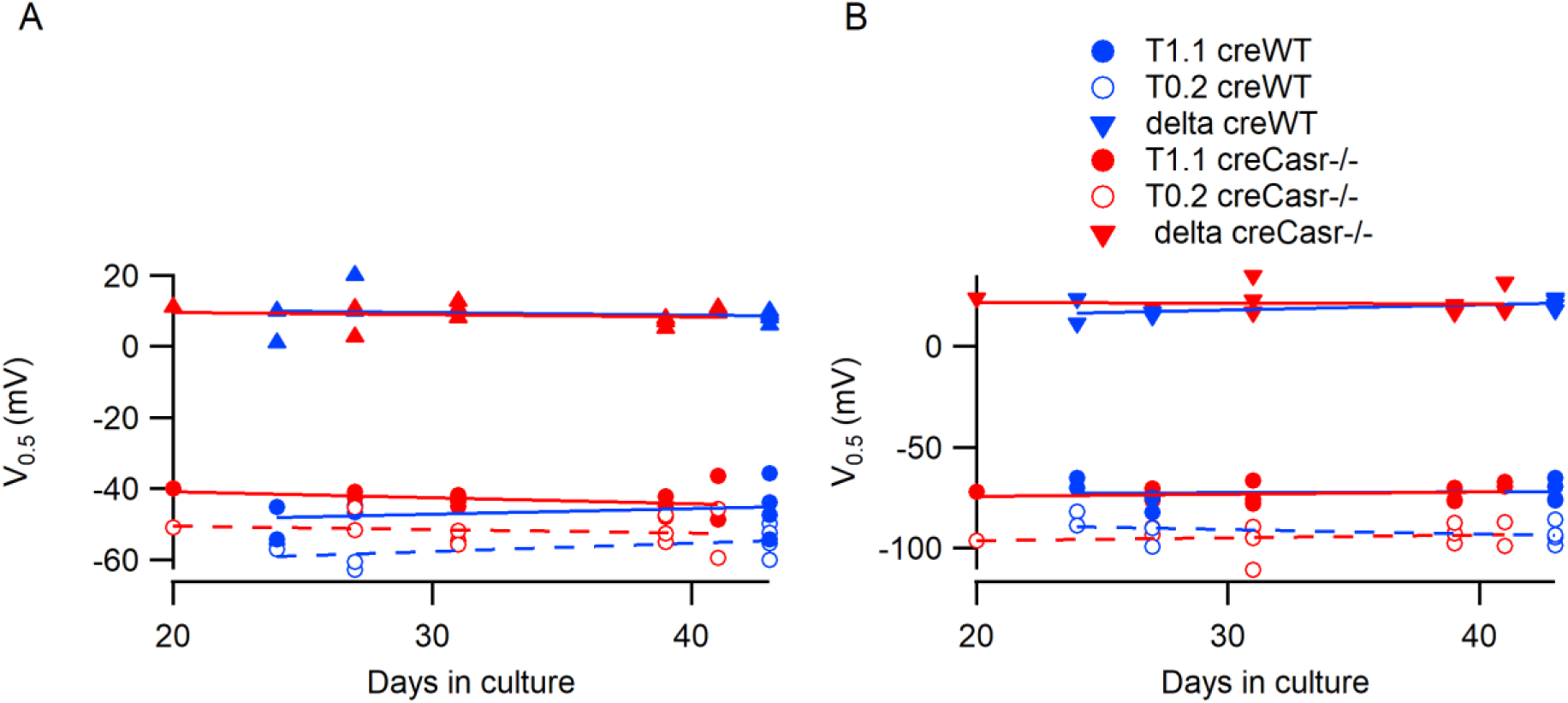
Culture age does not impact V_0.5_ for VGSC current activation (A) and inactivation (B) in patches from creWT and creCasr-/- neurons. The delta (triangles) represents the shift in V_0.5_ fllowing the switch from T_1.1_ to T_0.2._

**Figure 5.**
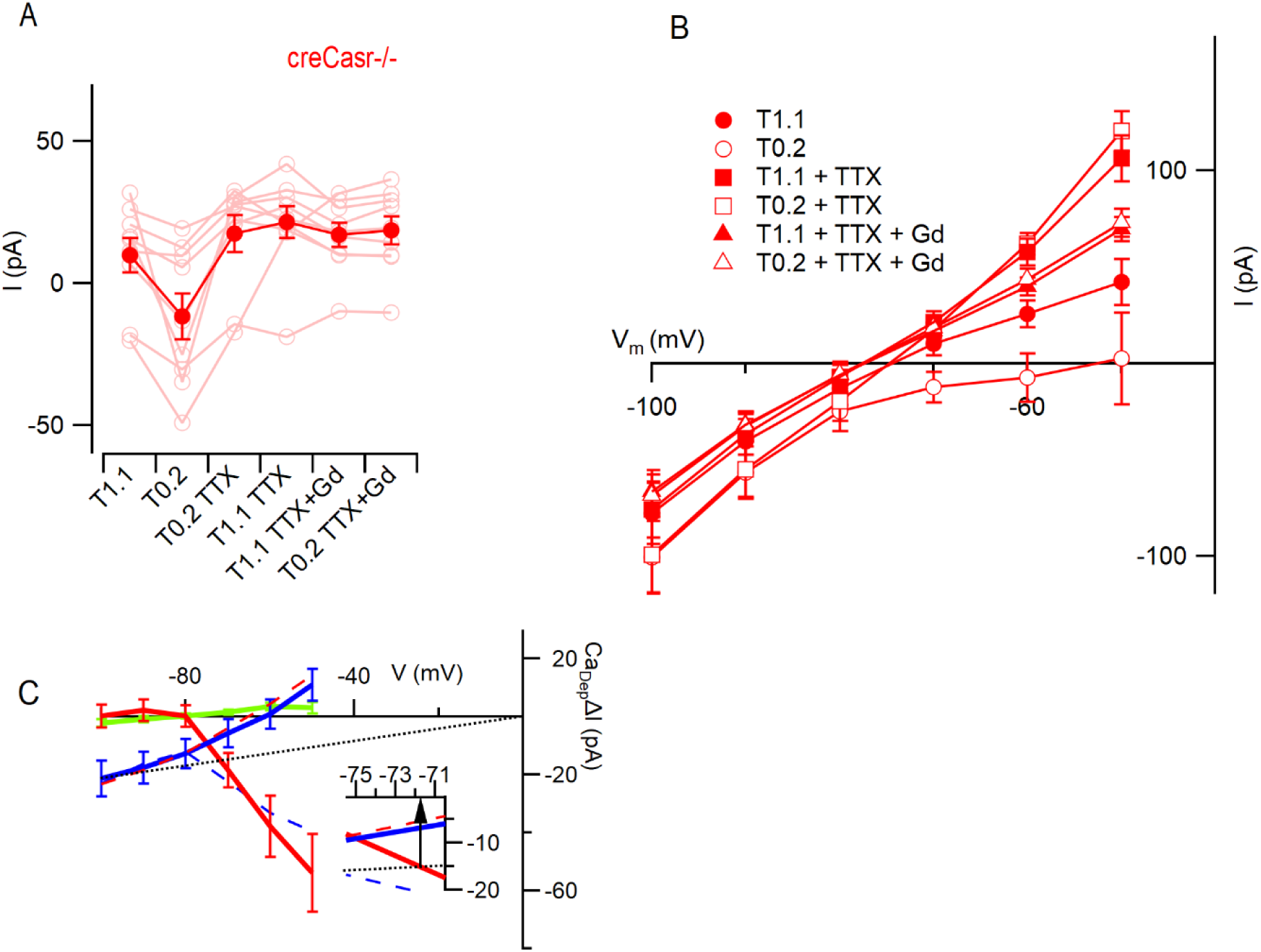
VGSC current activation by decreased [Ca^2+^]_o_. Supp Figure A) Plot of average basal current measurements (filled circles) and individual neurons (open circles) in each solution condition in creCasr-/- (n = 9) neurons. Each basal current represents the average value recorded during last 20 s of the specific solution application. B) Plots of average currents versus voltage for creCasr-/- (n = 9) neurons as per Figure 5 F C) Plots of the average [Ca^2+^]_o_ dependent currents derived by subtraction of creCasr-/- data as per Figure 5 G.

## Supplementary Data

**ANOVA and post-hoc analysis statistical tables**

**Supplementary Table 1.** Action potential frequency.

| ANOVA table | SS | DF | MS | F (DFn, DFd) | P value |
| --- | --- | --- | --- | --- | --- |
| Interaction | 130.0 | 2 | 64.99 | F (2, 54) = 3.193 | P = 0.0489 |
| [Ca<sup>2+</sup>]<sub>o</sub> on AP count | 594.4 | 1 | 594.4 | F (1, 54) = 29.21 | P < 0.0001 |
| Genotype | 136.1 | 2 | 68.04 | F (2, 54) = 3.368 | P = 0.0418 |
| Subjects (matching) | 1091 | 54 | 20.20 | F (54, 54) = 0.9925 | P = 0.5110 |
| Residual | 1099 | 54 | 20.35 |  |  |

**Supplementary Table 2.** Action potential frequency.

| ANOVA table | SS | DF | MS | F (DFn, DFd) | P value |
| --- | --- | --- | --- | --- | --- |
| Interaction | 0.3407 | 1 | 0.3407 | F (1, 35) = 0.4758 | P = 0.4949 |
| [Ca<sup>2+</sup>]<sub>o</sub> at hyperpolarizing injection | 8.262 | 1 | 8.262 | F (1, 35) = 11.54 | P = 0.0017 |
| Genotype | 0.5380 | 1 | 0.5380 | F (1, 35) = 0.7309 | P = 0.3984 |
| Subjects (matching) | 25.76 | 35 | 0.7360 | F (35, 35) = 1.028 | P = 0.4679 |
| Residual | 25.06 | 35 | 0.7161 |  |  |

**Supplementary Table 3.** Action potential frequency.

| ANOVA table | SS | DF | MS | F (DFn, DFd) | P value |
| --- | --- | --- | --- | --- | --- |
| Interaction | 0.6090 | 1 | 0.6090 | F (1, 35) = 0.2048 | P = 0.6536 |
| [Ca<sup>2+</sup>]<sub>o</sub> at depolarizing injection | 45.09 | 1 | 45.09 | F (1, 35) = 15.17 | P = 0.0004 |
| genotype | 5.982 | 1 | 5.982 | F (1, 35) = 0.9959 | P = 0.3252 |
| Subjects (matching) | 210.2 | 35 | 6.006 | F (35, 35) = 2.020 | P = 0.0204 |
| Residual | 104.1 | 35 | 2.973 |  |  |

**Supplementary Table 4.** Action potential threshold.

| ANOVA table | SS | DF | MS | F (DFn, DFd) | P value |
| --- | --- | --- | --- | --- | --- |
| Interaction | 0.5225 | 1 | 0.5225 | F (1, 27) = 0.07478 | P = 0.7866 |
| [Ca<sup>2+</sup>]<sub>o</sub> on AP threshold | 394.6 | 1 | 394.6 | F (1, 27) = 56.48 | P < 0.0001 |
| Genotype | 54.34 | 1 | 54.34 | F (1, 27) = 2.284 | P = 0.1424 |
| Subjects (matching) | 642.5 | 27 | 23.80 | F (27, 27) = 3.406 | P = 0.0011 |
| Residual | 188.6 | 27 | 6.987 |  |  |

**Supplementary Table 5.** RMP.

| ANOVA table | SS | DF | MS | F (DFn, DFd) | P value |
| --- | --- | --- | --- | --- | --- |
| Interaction | 14.36 | 1 | 14.36 | F (1, 37) = 1.035 | P = 0.3155 |
| [Ca<sup>2+</sup>]<sub>o</sub> on RMP | 438.9 | 1 | 438.9 | F (1, 37) = 31.65 | P < 0.0001 |
| Genotype | 1513 | 1 | 1513 | F (1, 37) = 19.10 | P < 0.0001 |
| Subjects (matching) | 2930 | 37 | 79.19 | F (37, 37) = 5.710 | P < 0.0001 |
| Residual | 513.2 | 37 | 13.87 |  |  |

**Supplementary Table 6.** Action potential frequency.

| ANOVA table | SS | DF | MS | F (DFn, DFd) | P value |
| --- | --- | --- | --- | --- | --- |
| Interaction | 0.4305 | 3 | 0.1435 | F (3, 87) = 0.3481 | P = 0.7906 |
| [Ca<sup>2+</sup>]<sub>o</sub> and I | 22.23 | 3 | 7.410 | F (3, 87) = 17.97 | P < 0.0001 |
| Genotype | 0.1341 | 1 | 0.1341 | F (1, 29) = 0.2005 | P = 0.6577 |
| Subjects (matching) | 19.41 | 29 | 0.6692 | F (29, 87) = 1.623 | P = 0.0445 |
| Residual | 35.87 | 87 | 0.4123 |  |  |

**Supplementary Table 7.** Action potential frequency.

| Sidak's multiple comparisons test | Mean Diff. | 95% CI of diff. | Significant? | Summary | Adjusted P Value |
| --- | --- | --- | --- | --- | --- |
| Ia vs. T<sub>0.2</sub> Ia | -1.059 | -1.498 to -0.6200 | Yes | **** | < 0.0001 |
| Ia vs. T<sub>0.2</sub> Ib | -0.6203 | -1.059 to -0.1813 | Yes | ** | 0.0016 |
| Ia vs. Ib | -0.1163 | -0.5554 to 0.3227 | No | ns | 0.9797 |
| T<sub>0.2</sub> Ia vs. T<sub>0.2</sub> Ib | 0.4387 | -0.0003709 to 0.8778 | No | ns | 0.0503 |
| T<sub>0.2</sub> Ia vs. Ib | 0.9427 | 0.5037 to 1.382 | Yes | **** | < 0.0001 |
| T<sub>0.2</sub> Ib vs. Ib | 0.5040 | 0.06496 to 0.9431 | Yes | * | 0.0160 |

**Supplementary Table 8.** Membrane potential with Ia.

| ANOVA table | SS | DF | MS | F (DFn, DFd) | P value |
| --- | --- | --- | --- | --- | --- |
| Interaction | 131.7 | 1 | 131.7 | F (1, 29) = 4.055 | P = 0.0534 |
| [Ca<sup>2+</sup>]<sub>o</sub> | 949.2 | 1 | 949.2 | F (1, 29) = 29.22 | P < 0.0001 |
| Genotype | 162.7 | 1 | 162.7 | F (1, 29) = 4.874 | P = 0.0353 |
| Subjects (matching) | 968.0 | 29 | 33.38 | F (29, 29) = 1.028 | P = 0.4711 |
| Residual | 942.0 | 29 | 32.48 |  |  |

**Supplementary Table 9A.**
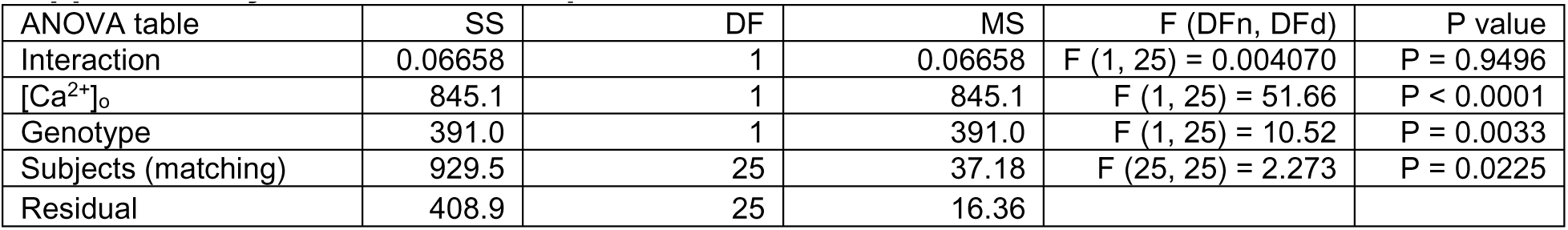
Action potential threshold recorded at −70 mV.

| ANOVA table | SS | DF | MS | F (DFn, DFd) | P value |
| --- | --- | --- | --- | --- | --- |
| Interaction | 0.06658 | 1 | 0.06658 | F (1, 25) = 0.004070 | P = 0.9496 |
| [Ca<sup>2+</sup>]<sub>o</sub> | 845.1 | 1 | 845.1 | F (1, 25) = 51.66 | P < 0.0001 |
| Genotype | 391.0 | 1 | 391.0 | F (1, 25) = 10.52 | P = 0.0033 |
| Subjects (matching) | 929.5 | 25 | 37.18 | F (25, 25) = 2.273 | P = 0.0225 |
| Residual | 408.9 | 25 | 16.36 |  |  |

**Supplementary Table 9B.** Action potential half-width recorded at −70 mV.

| ANOVA table | SS | DF | MS | F (DFn, DFd) | P value |
| --- | --- | --- | --- | --- | --- |
| Interaction | 2.008e-07 | 1 | 2.008e-07 | F (1, 28) = 2.800 | P = 0.1054 |
| [Ca<sup>2+</sup>]<sub>o</sub> | 1.415e-06 | 1 | 1.415e-06 | F (1, 28) = 19.73 | P = 0.0001 |
| Genotype | 3.050e-07 | 1 | 3.050e-07 | F (1, 28) = 0.4545 | P = 0.5057 |
| Subjects (matching) | 1.879e-05 | 28 | 6.710e-07 | F (28, 28) = 9.358 | P < 0.0001 |
| Residual | 2.008e-06 | 28 | 7.170e-08 |  |  |

**Supplementary Table 9C.** Action potential peak voltage recorded at −70 mV.

| ANOVA table | SS | DF | MS | F (DFn, DFd) | P value |
| --- | --- | --- | --- | --- | --- |
| Interaction | 0.0001602 | 1 | 0.0001602 | F (1, 28) = 6.758 | P = 0.0147 |
| [Ca<sup>2+</sup>]<sub>o</sub> | 0.0004821 | 1 | 0.0004821 | F (1, 28) = 20.34 | P = 0.0001 |
| Genotype | 0.001193 | 1 | 0.001193 | F (1, 28) = 5.891 | P = 0.0219 |
| Subjects (matching) | 0.005669 | 28 | 0.0002025 | F (28, 28) = 8.541 | P < 0.0001 |
| Residual | 0.0006637 | 28 | 2.370e-005 |  |  |

**Supplementary Table 10.** Voltage-gated sodium channel current V_0.5_ for inactivation.

| ANOVA table | SS | DF | MS | F (DFn, DFd) | P value |
| --- | --- | --- | --- | --- | --- |
| Interaction | 12.00 | 1 | 12.00 | F (1, 18) = 0.7743 | P = 0.3905 |
| [Ca<sup>2+</sup>]<sub>o</sub> | 3973 | 1 | 3973 | F (1, 18) = 256.2 | P < 0.0001 |
| Genotype | 27.49 | 1 | 27.49 | F (1, 18) = 0.5632 | P = 0.4627 |
| Subjects (matching) | 878.5 | 18 | 48.81 | F (18, 18) = 3.148 | P = 0.0097 |
| Residual | 279.1 | 18 | 15.50 |  |  |

**Supplementary Table 11.**
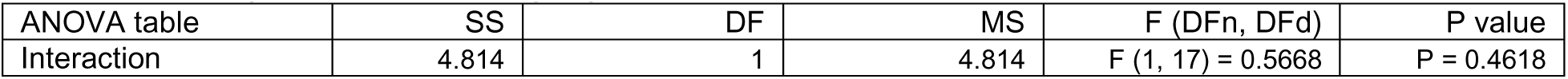
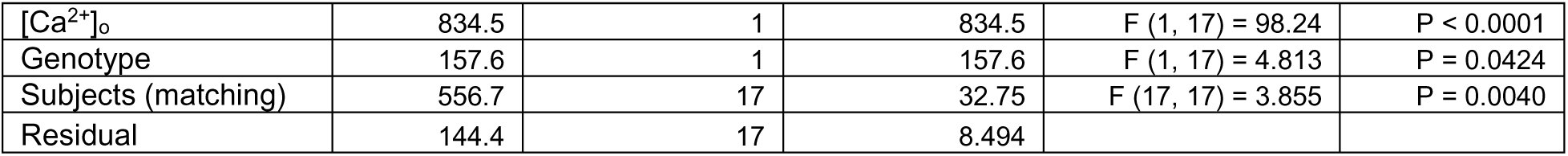
Voltage-gated sodium channel current V_0.5_ for activation.

**Supplementary Table 12.** Voltage-gated potassium channel currents at 60 mV.

| ANOVA table | SS | DF | MS | F (DFn, DFd) | P value |
| --- | --- | --- | --- | --- | --- |
| Interaction | 1.226e-018 | 3 | 4.086e-019 | F (3, 57) = 0.2271 | P = 0.8772 |
| [Ca<sup>2+</sup>]<sub>o</sub> and time | 7.270e-018 | 3 | 2.423e-018 | F (3, 57) = 1.347 | P = 0.2683 |
| Genotype | 6.054e-017 | 1 | 6.054e-017 | F (1, 19) = 1.231 | P = 0.2811 |
| Subjects (matching) | 9.345e-016 | 19 | 4.919e-017 | F (19, 57) = 27.33 | P < 0.0001 |
| Residual | 1.026e-016 | 57 | 1.800e-018 |  |  |

**Supplementary Table 13.**
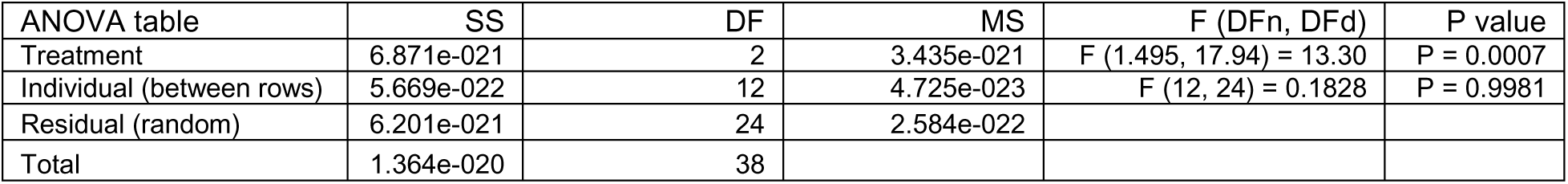
[Ca^2+^]_o_-dependent basal currents at −70 mV.

**Supplementary Table 14.** Post hoc testing of [Ca^2+^]_o_-dependent basal currents at −70 mV.

| Sidak's multiple comparisons test | Mean Diff. | 95% CI of diff. | Significant? | Summary | Adjusted P Value |
| --- | --- | --- | --- | --- | --- |
| TTX sens vs. Gd sens | -2.247e-011 | -4.391e-011 to -1.033e-012 | Yes | * | 0.0392 |
| TTX sens vs. Rem | -3.158e-011 | -4.899e-011 to -1.418e-011 | Yes | *** | 0.0009 |
| Gd sens vs. Rem | -9.111e-012 | -2.146e-011 to 3.240e-012 | No | ns | 0.1789 |

**Supplementary Table 15.**
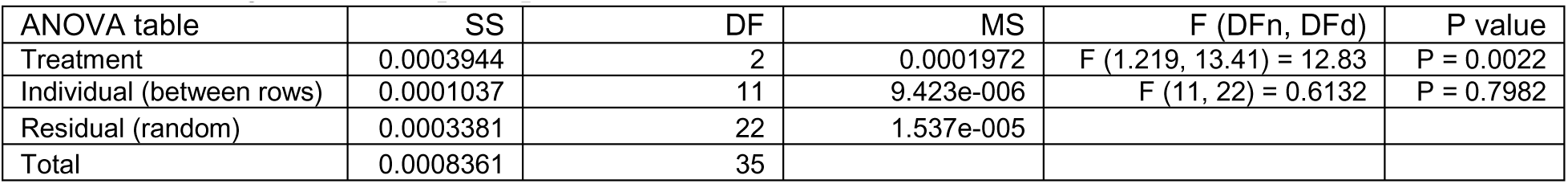
[Ca^2+^]_o_-dependent depolarization.

**Supplementary Table 16.** Post hoc testing of blocker sensitive fractions of the [Ca^2+^]_o_-dependent depolarization.

| Sidak's multiple comparisons test | Mean Diff. | 95% CI of diff. | Significant? | Summary | Adjusted P Value |
| --- | --- | --- | --- | --- | --- |
| TTX sens vs. Gd sens | 0.006528 | 0.001005 to 0.01205 | Yes | * | 0.0215 |
| TTX sens vs. Rem | 0.007428 | 0.002879 to 0.01198 | Yes | ** | 0.0028 |
| Gd sens vs. Rem | 0.0009 | -0.001301 to 0.003101 | No | ns | 0.5311 |

